# Exploring Probiotic Potential: A Comparative Genomics and In Silico Assessment of Genes within the Genus *Geobacillus*

**DOI:** 10.1101/2024.05.15.594408

**Authors:** Ishfaq Nabi Najar, Prayatna Sharma, Rohit Das, Krishnendu Mondal, Ashish Kumar Singh, Anu Radha, Varsha Sharma, Sonali Sharma, Nagendra Thakur, Sumit G. Gandhi, Vinod Kumar

**Affiliations:** Fermentation and Microbial Biotechnology Division, CSIR IIIM, Jammu India; Department of Microbiology, School of Life Sciences, Sikkim University, 6th Mile, Samdur, Tadong, Gangtok – 737102, Sikkim, India; Department of Microbiology, Vidyasagar University, Midnapur India; Department of Biotechnology and Synthetic Biology, CIAB, Mohali, India; Infectious Diseases Division. CSIR IIIM Jammu, India; Fermentation and Microbial Biotechnology Division, CSIR IIIM, Jammu India.

**Keywords:** *Geobacillus*, Probiotics, Pan-Genome, Comparative Genomics, Probiogenomics, Phylogenetics

## Abstract

The pursuit of new probiotic targets has seen a surge, aided by next-generation sequencing, facilitating a thorough exploration of bacterial traits. The genus *Geobacillus* stands out as a promising target for uncovering its potential as a probiotic. The study explored the genetic attributes of the genus *Geobacillus* for their resilience to gastrointestinal conditions, nutrient production, and immunomodulatory compound creation, revealing potential probiotic traits. Additionally, the research undertook predictive analyses of genomic elements such as prophages, CRISPR-Cas systems, insertion sequences, genomic islands, antibiotic resistance genes, and CAZymes. These evaluations aimed to assess the safety aspects associated with the genus *Geobacillus*. A comparative genomic analysis was also carried out using 18 validly published genomes of the genus *Geobacillus* and a few other genomes of *Lactobacillus* and *Bifidobacterium* were taken as control. Genes associated with probiotic traits like adhesion, stress tolerance (acid/bile, osmotic, oxidative), immune modulation, and molecular chaperones were uniformly detected in the *Geobacillus* genus. Notably, mobile genetic elements such as plasmids, prophages, and insertion sequences were absent, as were virulence factors, toxins, and Antibiotic resistance genes. Additionally, CRISPR-Cas systems and CAZymes were present. The pan-genome encompassed 25,284 protein-coding genes with translation. Comparative genomic analysis revealed an open pan-genome for *Geobacillus*. Pan-genome exhibited variability, particularly in genes linked to environmental interaction and secondary metabolite synthesis. In conclusion, *Geobacillus* appears potentially safe and well-suited for the gut habitat. However, further *in vitro* studies are essential to add to the knowledge of the probiotic potential of *Geobacillus* species.

**Importance:** This comprehensive study highlights the significant probiotic potential and genetic makeup of the *Geobacillus* genus, shedding light on its unique attributes in adapting to extreme environmental conditions. Understanding the probiotic properties of *Geobacillus* is crucial amidst growing concerns over antibiotic resistance, offering promising alternatives for combating pathogenic microbes. Additionally, exploring the genetic diversity and adaptive mechanisms of *Geobacillus* through genomic and metagenomic approaches provides valuable insights into its biotechnological applications and evolutionary history. By employing in-silico methods and comparative analyses with established probiotic genera, this study contributes to elucidating the probiotic characteristics of *Geobacillus*, paving the way for further research in harnessing its beneficial traits for various applications in health, biotechnology, and environmental remediation.

## Introduction

The Earth encompasses unique repositories characterized by extreme environmental conditions, where some microbes termed extremophiles thrive with the aid of evolutionary adaptations required for survival (1). Particularly, the genus *Geobacillus* belongs to obligate thermophilic, Gram-positive rods, aerobic or facultative aerobic species, thriving in the warmest regions on earth, from geothermal springs to equatorial deserts. Surviving in regions accustomed to 45-80℃, these obligatory thermophiles possess optimum growth at 55-65℃ (2, 3). Previously known to belong to the genus *Bacillus*, the reclassified genus *Geobacillus* holds immense biotechnological importance. Their habitat remains remarkably diverse, ranging from cold environments such as Antarctica (4) to extremely hot niches (5). Their ability to withstand harsh temperature conditions paves their way towards numerous biotechnological applications; these species serve as key hosts for thermostable proteins with robust catalytic activity, such as amylases, protease, and lipases; and also serve various roles in bioprocessing applications such as bioproduction of biofuels and biodiesel and bioremediation (2, 6).

The genus *Geobacillus* was previously classified as a separate unit, known as group 5 of the thermophilic genus *Bacillus*, as demonstrated by a 16S rRNA gene sequencing study by Ash et al, 1991 (7). However, due to genetic heterogeneity, the revision of taxonomic classification let the group be reclassified as *Geobacillus* (7). Currently, there are 20 species prescribed under the genus *Geobacillus* (8). Research since the last century has involved the reclassification of the genus *Bacillus* to *Geobacillus*, with the earliest report of isolation of *Bacillus stearothermophilus* (now termed *Geobacillus stearothermophilus*) by Donk (9). Until the end of the 21^st^ century, a taxonomic study carried out by (10) revealed species of *B. kaustophilus*, *B. stearothermophilus*, *B. thermocatenulatus*, *B. thermodenitrificans*, *B. thermoglucosidasius*, and *B. thermoleovorans* exhibited similar physiological and phylogenetic properties, thereby classifying these species into a new genus “*Geobacillus*”. With most of the species with a Gram-positive nature and few showing a Gram-variable nature, the bacteria appear in short chains and may or may not be motile. With an optimum temperature for growth at 55–65 °C, these species are obligate thermophiles and exhibit growth in neutral pH ranges (6.0–8.5) (1). The nutritional profile of *Geobacillus* emerges as chemoorganotrophs, with oxygen as the terminal electron acceptor (with few cases showing nitrate as an electron acceptor). The species require no special nutrients like vitamins and can be grown in simple media such as nutrient agar, with basic requirements like carbohydrates, peptone, and tryptone as primary carbon and energy sources (11). Various studies have been done to evaluate the biotechnological potential of the *Geobacillus* species, however, there are very few studies done on the probiotic potential of the genus *Geobacillus*.

Probiotics that are often considered live microbes confer health benefits when administered due to their antagonistic activity against their surrounding microbes (12). Considering the immense global threat that antibiotic resistance poses to humanity today (13), there remains a constant need for novel strains with enhanced probiotic properties that confer stronger activity against pathogenic microbes. Members belonging to the *Firmicutes* have gained considerable attention for their probiotic properties for humans and animals (14, 15). The genera *Geobacillus* and *Parageobacillus* secrete anti-microbial compounds such as bacteriocins, antibiotics, and bacteriocin-like inhibitory substances (BLISes) and serve as potential candidates exhibiting unique probiotic properties (12). For instance, a preliminary study on *Geobacillus thermoleovorans* isolated from oil waste showed antimicrobial activity against pathogenic strains of *Salmonella typhimurium*, *Vibrio parahemolitycu*s, *Staphylococcus aureus* as evident by antagonism and adherence assay, denoting the strain as a potential probiotic candidate (16). The ability to adhere to abiotic surfaces to create a bio-secure environment may assist in inhibiting the growth of pathogenic strains (12). Other mechanisms include the inhibition of Quorum-sensing in Gram-negative bacteria by *Geobacillus* strains, due to their Acyl Homoserine Lactones (AHLs) degrading ability (17). The plausible explanation for their candidature in the ever-increasing probiotic market may be due to their enhanced survival strategies as compared to the routinely used probiotic strains. Additionally, these organisms are part of a group that is both biologically safe and non-pathogenic. They are recognized as a bimodal probiotic microbiota present in both humans and animals and are an essential composite of the soil, skin, gastrointestinal, and plant microbiome (15). The strains of *Geobacillus* thrive in extreme environmental parameters of high temperature, different pH ranges, UV radiation, heavy metals, salts, organic/inorganic chemicals and detergents, and arid and minuscule moisture environments, mainly owned by their ability to form durable endospores (12, 15).

Various reports suggest the prevalence of antibiotic resistance in mesophilic bacterial species, however, the phenomenon of antibiotic resistance in thermophiles is lesser known. It remains crucial to unravel the antibiotic resistance patterns and the intricate mechanism involved in thermophilic bacteria, given the immense biotechnological applications that the thermophilic bacteria impose (Najar et al., 2020c). Understanding antibiotic resistance in thermophiles remains crucial. Very few studies suggest the study of antibiotic-resistance profiling among thermophilic bacteria. Given this, Najar et al, 2022 conducted research to study antibiotic resistance prevailing in the microbiota of Sikkim hot springs. Both the culture-dependent and PCR amplification analyses revealed the absence of antibiotic resistance by the thermophilic bacteria. Similarly, no antibiotic-resistance genes were found in *Geobacillus* species, suggesting that the genus susceptible to antibiotics natively (13). However, it is crucial to evaluate the genetic makeup of these *Geobacillus* species, particularly emphasizing the diverse attributes that contribute to their probiotic potential using metagenomic and pan-genomic approaches.

One of the approaches such as standard Sanger sequencing is well known and over the years, numerous sequencing platforms like the next generation sequencing have emerged that employ different sequencing strategies to unravel the genomic and metabolic potential of the genome (19). Genome sequencing infers to unraveling the sequences of the entire genome of an organism, instead of sequencing individual genes (20). The whole genome shotgun sequencing method helps in understanding the metabolic and genomic potential of the repertoire of genes present in an organism. Deciphering the complete genome of an organism helps to provide insight into understanding the metabolic potential, metabolomics, functional genes, and adaptive mechanisms (21). For instance, the whole genome sequence of a putative novel strain of thermophilic bacterium *Parageobacillus* sp. indicates the presence of various important genes for carbohydrate, Sulphur, nitrogen, and phosphorus metabolism (21). Another widely employed method is the metagenomic approach which helps in elucidating the diversity of microbes present and crucial genes that are responsible for conferring adaptive strategies to the microbiome, such as antibiotic resistance or stress-related genes. Functional metagenomic study on hot spring samples of Sikkim hot springs reveal diverse set of genes required to cope with stressful conditions such as heat shock, acid, and osmotic stress; various multidrug efflux pumps and transport systems, genes for degradation of xenobiotics, etc. (22).

Pan-genome analyses are a powerful tool in bacterial taxonomy that may help redefine the categorization of species or, in defining new species (23). The term pan-genome represents the entire set of genes present in a species and is categorized into three main categories-a core genome that comprises sequences shared between all the species; an accessory genome that represents genes shared in some species; and a singleton or unique genome that is present in a particular species only. The core genome comprises genes required for basic metabolic activities such as genes for antibiotic resistance and housekeeping genes. These genes that are conserved usually infer evolutionary relationships amongst different strains. The genes present in the accessory genome participate in adaptive strategies in a strain, where horizontal gene transfer (thus are subjected to gene gain/loss) plays a major role that aids in coping up adaptive mechanisms in a bacterium when subjected to a new niche (24). Pan-genome analysis plays a major role in implicating the evolution of the genus *Geobacillus* from *Bacillus.* There remains limited study about pan-genome analyses of the *Geobacillus* genus, although reports for other bacteria from the neighboring genus *Bacillus* have been reported (25). Bezuidt et al, 2016, provided insights into the role of horizontal gene transfer in the diversification of *Geobacillus* (26).

This study employs an in-silico approach to assess the potential probiotic characteristics of the Geobacillus genus. It delves into various attributes, including taxonomy, phylogenomics, probiotic-related genes, prediction of antibiotic resistance genes, virulence factors, toxins, mobile genetic elements, plasmids, bacteriocins, CRISPR-Cas systems, and CAZymes. Additionally, a pan-genome analysis was conducted. To benchmark and contextualize the findings, comparative analysis involved two established genera, *Bifidobacterium* and *Lactobacillus*, serving as positive controls. Overall, this comprehensive study serves as a significant exploration of both the probiotic potential and genetic makeup of the *Geobacillus* genus.

## Results

### Source and quality check of various genomes

Among 18 *Geobacillus* species used in this study, 13 are validly published under ICNP as per LPSN and among 13 *Geobacillus* species used, 7 are type species such as *Geobacillus icigianus* G1w1, *Geobacillus kaustophilus* NBRC 102445, *Geobacillus lituanicus* N-3, *Geobacillus proteiniphilus* 1017, *Geobacillus stearothermophilus* ATCC 12980, *Geobacillus thermocatenulatus* BGSC 93A1, and *Geobacillus thermodenitrificans* DSM 465. In the case of control, probiotic species used are validly published and type species. The quality of genomes was assessed and it was shown that all the genomes possess good quality with >97% completeness and very low fractional contamination as shown in Table 1**. and Supplementary Report File.1.** The GC content of *Geobacillus* genomes range between 49-53%, *Lactobacillus* genomes range between 34-49% and to that of *Bifidobacterium* genomes range between 58-60%. The total genome length of the *Geobacillus* genomes was higher (3.45mb-3.80mb) than those of *Lactobacillus* (2.03mb-3.2mb), and *Bifidobacterium* genomes (2.23mb-2.83mb). The isolation source of most of the *Geobacillus* species analyzed in this study is environmental such as Oil fields, Hot springs, fumaroles, and sediments. However, the isolation source of Geobacillus *stearothermophilus* ATCC 12980 and *Geobacillus thermodenitrificans* DSM 465 was food products such as spoiled canned food and sugar beet juice respectively. On the other hand, the isolation source of other probiotic species was food and gastrointestinal tract as shown in **Table 1**.

**Table 1.** Quality control, source and GC content of studied genomes.

| Genomes | Source | Quast |  |  | CheckM |  |
| --- | --- | --- | --- | --- | --- | --- |
|  |  | Total Length | N50 | GC | Completeness | Contamination |
| *Geobacillus_icigianus_G1w1_NZ_JPYA01000193* | Hot spring sediments | 3457810 | 39998 | 52.03 | 99.45 | 0.34 |
| *Geobacillus_kaustophilus_NBRC_102445_NZ_BBJV01000028* | Pasteurized milk | 3454242 | 50413 | 51.86 | 98.31 | 0 |
| *Geobacillus_jurassicus_NBRC_107829_BCQG0000000* | Unknown | 3670957 | 3670957 | 52.22 | 99.45 | 0 |
| *Geobacillus_lituanicus_N-3_CP017692* | High-temperature oilfield | 3499511 | 3447958 | 52.14 | 98.91 | 0.55 |
| *Geobacillus_proteiniphilus_1017_NZ_MQMG0000000* | High-temperature heavy oil reservoir | 3574950 | 52882 | 51.76 | 99.45 | 0.68 |
| *Geobacillus_stearothermophilus_ATCC_12980_NZ_JYNW01000038* | Spoiled canned food | 2860911 | 2831741 | 52.77 | 98.61 | 0 |
| *Geobacillus_subterraneus_KCTC3922_NZ_CP014342_1* | Oil field | 3474426 | 3474426 | 52.2 | 99.45 | 0 |
| *Geobacillus_thermocatenulatus_BGSC_93A1_N_EWK01000001* | Oil field | 3563800 | 1841649 | 51.77 | 99.45 | 0.03 |
| *Geobacillus_thermodenitrificans_DSM_465_NZ_AYKT01000065* | Sugar beet juice | 3400546 | 264274 | 49.05 | 99.45 | 0 |
| *Geobacillus_thermoleovorans_SURF-48B_NZ_NAGD00000000* | Yates Shaft, SURF | 3808577 | 3808577 | 52.19 | 99.45 | 1.5 |
| *Geobacillus_uzenensis_BGSC_92A1_NZ_NEWL00000000* | Oil field | 3357711 | 429603 | 52.23 | 99.45 | 0 |
| *Geobacillus_sp._GHH01_NC_020210* | Botanischer Garten | 3583134 | 3583134 | 52.28 | 99.45 | 0 |
| *Geobacillus_vulcani_PSS1_NZ_JPOI01000001* | Unknown | 3389115 | 3389115 | 52.4 | 99.45 | 1.39 |
| *Geobacillus_sp._CAMR5420_NZ_JHUS00000000* | Unknown | 3499823 | 95605 | 51.89 | 99.45 | 0.14 |
| *Geobacillus_sp._Y412MC52_CP002442* | hot spring | 3673940 | 3628883 | 52.34 | 99.45 | 0 |
| *Geobacillus_sp._C56-T3_NC_014206* | hot spring | 3650813 | 3650813 | 52.49 | 99.45 | 0 |
| *Geobacillus_sp._LEMMJ02_NZ_VKJO00000000* | Fumarole sediment | 3436274 | 150524 | 52.48 | 99.45 | 0.69 |
| *Geobacillus_zalihae*_JS1-2_CP129217 | Compost | 3567837 | 3532984 | 52.26 | 99.45 | 1.41 |
| *Lactobacillus_acidophilus*_ATCC_9224__NZ_CP130437 | Gastrointestinal tract | 2035536 | 2035536 | 34.76 | 98.3 | 0 |
| *Lactobacillus_delbrueckii*_subsp._*lactis*_DSM_20072_CP022988 | Emmental cheese | 2165984 | 2165984 | 49.05 | 100 | 0.75 |
| *Lactiplantibacillus_plantarum*_DSM_20174__CP039121 | Pickled cabbage | 3250154 | 3242936 | 44.5 | 99.07 | 2.78 |
| *Bifidobacterium_breve*_DSM_20213__BBAO00000 | Gastrointestinal tract | 2232812 | 35183 | 58.69 | 100 | 0 |
| *Bifidobacterium_longum*_subsp._*infantis*_ATCC_15697_CP001095 | Intestine of infant | 2832748 | 2832748 | 59.86 | 100 | 0 |

### Taxonomy and Phylogenomics Analysis

The ANI values were plotted on a heat map **Fig.1**. It has been shown that the *Geobacillus* genomes possess ANI values between 90-100%, thus predicting their high nucleotide identity. Whereas the *Lactobacillus* and *Bifidobacterium* genomes possess ANI values between 60-70%, also concerning genomes of genus *Geobacillus*, thus indicating their low nucleotide identity. The cluster analysis represented by the dendrogram shown in the heat map formed two major clades bifurcating the genus *Geobacillus* and two other genera studied i.e. *Lactobacillus* and *Bifidobacterium* based on ANI values. This supports the above assumption indicating that genomes of genus *Geobacillus* possess high nucleotide identity. The genus *Geobacillus* clade further differentiates into two main sub-clades. It was interesting to see that the earlier isolated and type species such as *G. stearothermophilus*, *G. icigianus*, *G. thermodenitrificans*, *G. lituanicus*, *G. uzenensis*, and *G. subterraneus* possess little variation in ANI values than the recently isolated genomes which are not validly published. The phylogenetic analysis among the genomes of only the genus *Geobacillus* shows a similar pattern with two major clades and three separate clades have been formed by *G. icigianus, G. jurassicus*, and *G. vulcani* **Fig.2**.

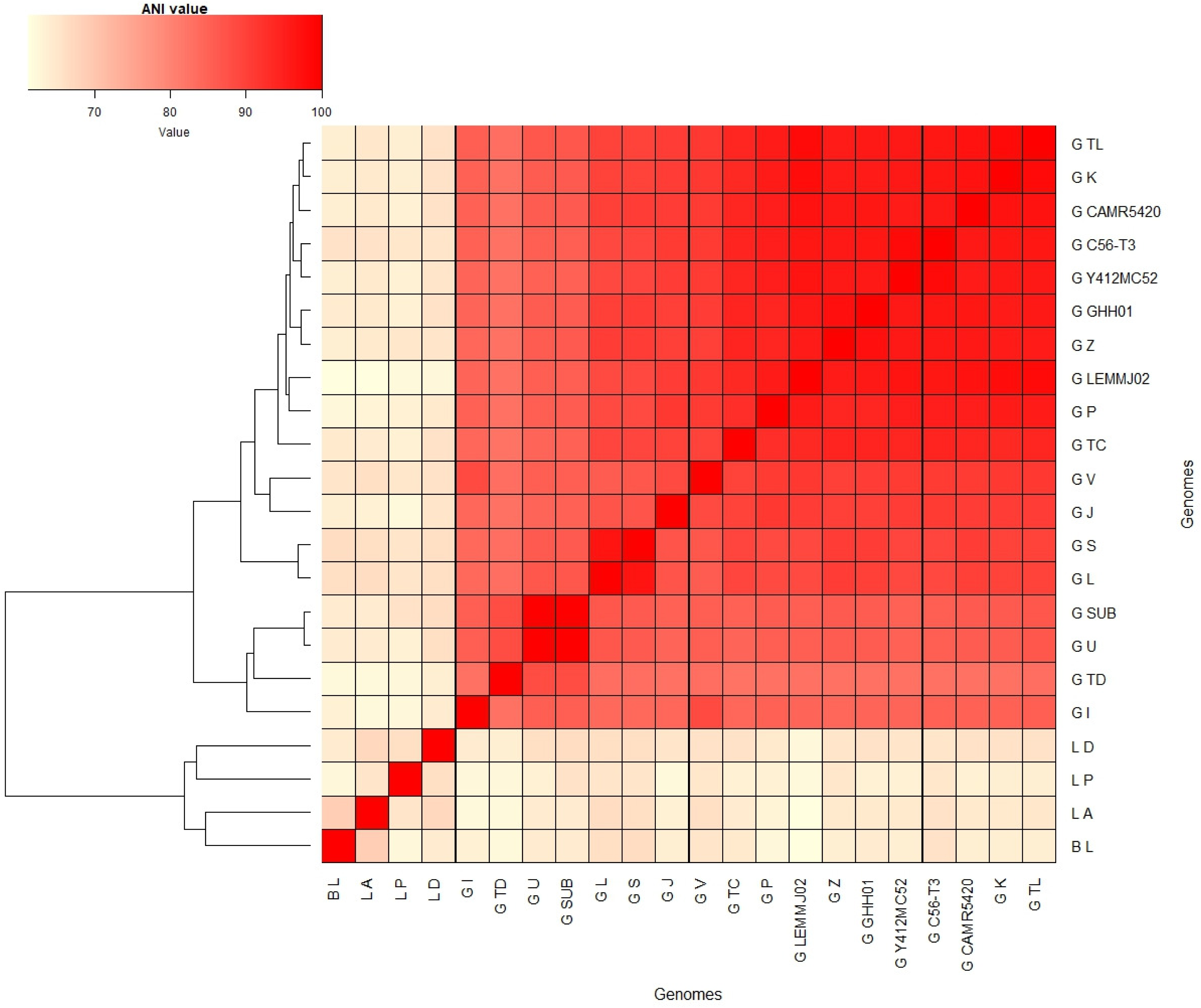

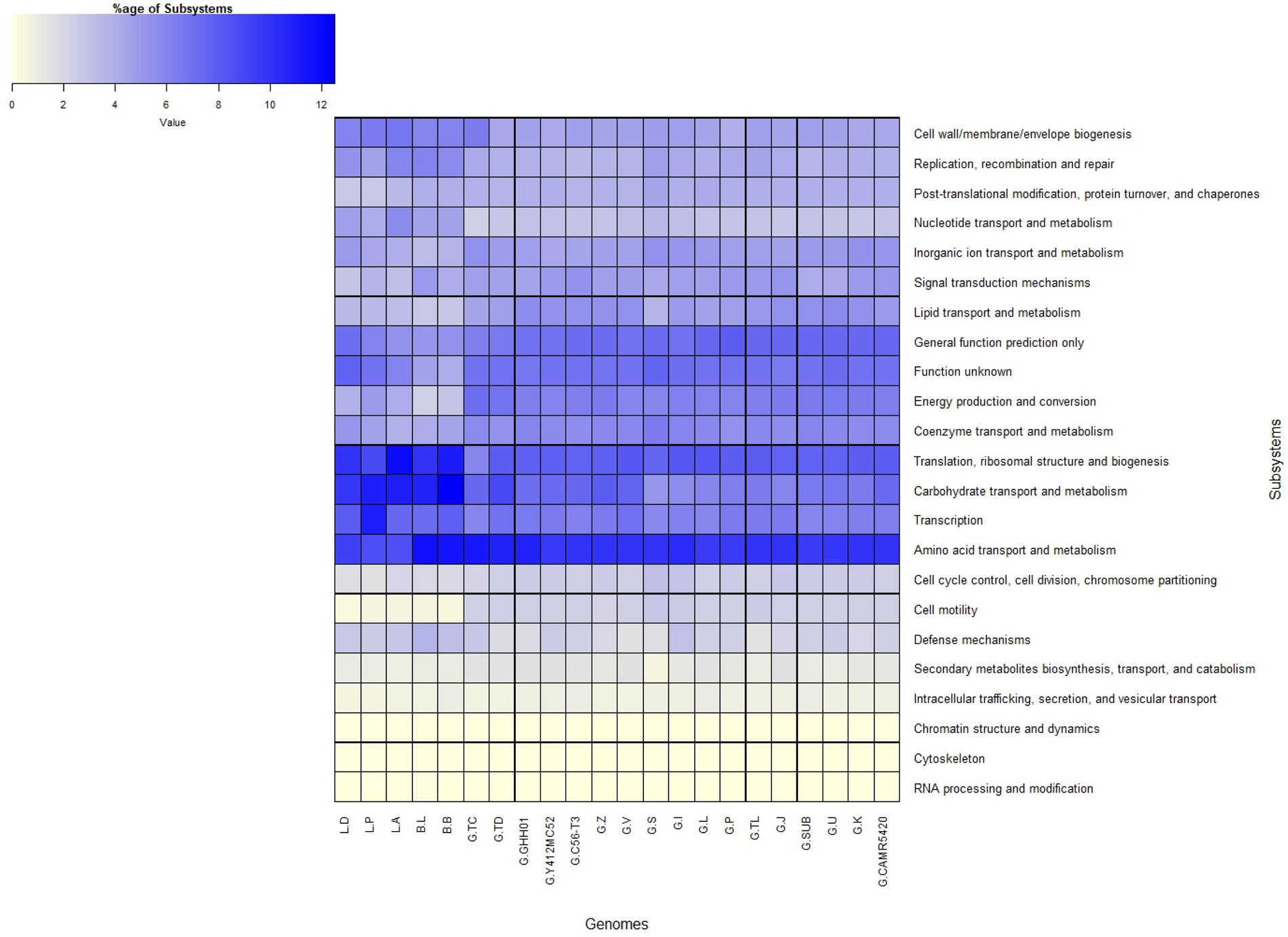

We were also trying to find out the phylogenetic relation among the genus *Geobacillus* and two other probiotic species of genus *Lactobacillus* and *Bifidobacterium* based on COG-based core genes **Fig.3**. and Supplementary Fig.1. and single copy genes using single-copy BV-BRC PGFams. The phylogenetic tree, established using a set of 49 core universal genes defined by COG, indicates the emergence of two primary clades encompassing the genus *Geobacillus* as well as the genus *Lactobacillus* and *Bifidobacterium*. The genus *Geobacillus* further differentiates into two sub clades, whereas species *G. jurassicus*, *G. vulcani*, and *G. icigianus form* separate clades similar to the phylogenetic tree of only genus *Geobacillus* based on ANI values. The genus *Lactobacillus* shows a close relation to *Parageobacillus* species whereas *Lactobacillus* and *Bifidobacterium* are distantly related to genus *Geobacillus* as they possess separate clades. Similar results were shown by phylogenetic tree based on single copy genes as shown in **Fig.4**. However, based on these genes *Lactobacillus* forms a clade close to *G. icigianus,* and other earlier isolated genomes such as *G. stearothermophilus*, *G. thermodenitrificans*, *G. lituanicus*, and *G. subterraneus.* Thus, based on core gene phylogenetic analysis of *Geobacillus*, *Lactobacillus,* and *Bifidobacterium* genomes, it has been shown that they significantly differ which is further supported by phylogenetic analysis based on single gene analysis.

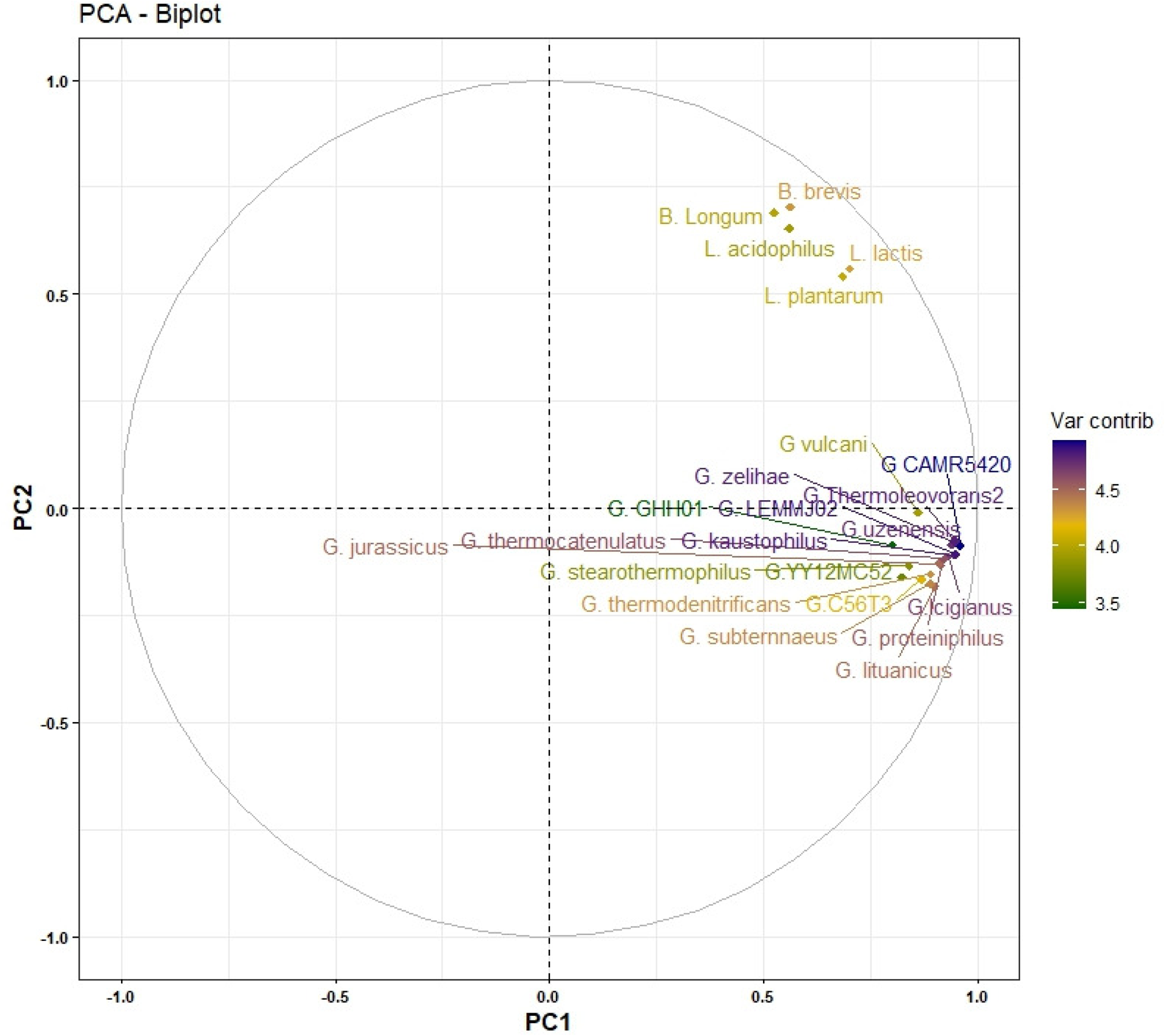

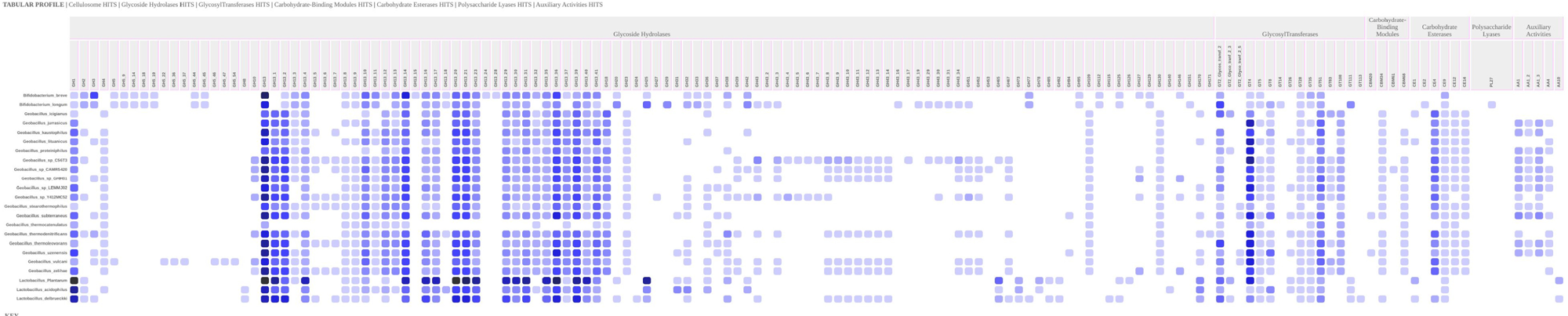

All the genomes were annotated with Prokka using the KBase platform. However, the cross-genus comparisons were also done using the new Comparative Systems service at BV-BRC which annotates the genomes based on global protein families, known as PGFams using RASTtk4. The functional assessment (subsystem analysis) among the genomes using canonical gene family assignment from COG, PFAM, TIGRFAM, and The SEED has been done using View Function Profile for Genomes - v1.4.0 and is represented in Heatmap **Fig.5**. it has been shown that subsystems like secondary metabolites biosynthesis, transport, catabolism, intracellular trafficking, secretion, chromatin structure and dynamics, cytoskeleton, and RNA processing and modification were uncharacterized in all the genomes. However, there is not much difference in other characterized subsystems between genomes of the genus *Geobacillus*, *Lactobacillus*, and *Bifidobacterium*. Cell motility is absent in the genus *Lactobacillus* and *Bifidobacterium* whereas present in the genus *Geobacillus*. Moreover, there are slight differences in a few subsystems among the studied genus such as Energy production and conversion, and lipid transport and metabolism. This signifies that there might be similarities among these genera considering probiotic properties.

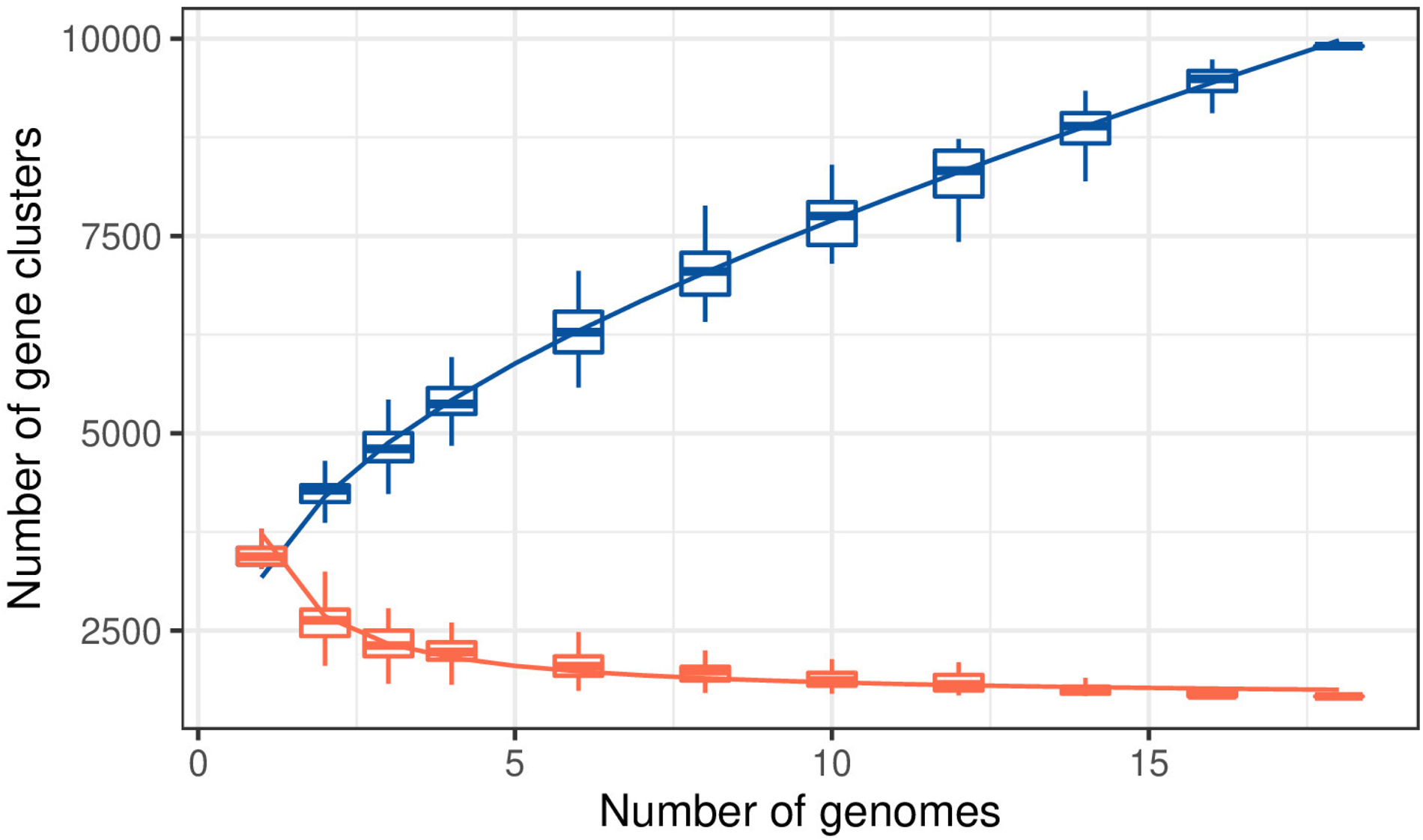

### Identification of Genes Related to Probiotics

Genes involved in the mechanism of modulation of the immune system, vitamin biosynthesis, fatty acid synthesis, resistance to stress conditions (acid, bile, osmotic, and oxidative stress), adhesion, bacterial colonization, and molecular chaperones were detected. The predicted genes among all these categories are present differently in **Table 2**. 25 genes of immune system modulation were screened and genes such as *Pyrc*, *PyrE*, *PyrH*, *PyrB*, *PyrG*, *PyrK*, *TxrA*, *TxrB*, *Clpb*, *CysC*, *nrdR* are present in all the genomes whereas *CysE*, *YjbK*, and *YjbM* were present only in *Geobacillus* genomes and *nrdH* and *dltC* are present only in *Bifidobacterium* and *Lactobacillus* genomes. 25 genes of adhesion were screened and genes such as *Tuf*, *clpB*, *clpX*, and *clpC* are present in all the genomes. *comC*, *srtD*, *PilT*, *PilZ*, and *lapA* are present in only the *Geobacillus* genomes whereas *dltD* and *dltA* were present in *Bifidobacterium* and *Lactobacillus* genomes only. Three sortase genes such as *strA1*, *strA2*, and *strA3* were absent in all genomes except *srtA1* was present in *Lactobacillus* genomes only. Also, *PilA*, *PilB*, *PilC*, *PilO*, and *PilM* adhesion genes were absent in all the genomes. Six bacterial colonization genes were screened, and only gene *TadA* was present in all the genomes rest were absent such as *TadB*, *TadC*, *TadE*, *TadF*, and *TadZ*. All the acid stress-related genes such as *atpB*, *atpC*, *atpD*, *atpE*, *atpF*, *atpG*, *atpH*, *recA*, *sodA*, *luxS*, *glmU*, *glmS*, *glmM1*, *glmM2* and asps were present in all the genomes whereas *cspA* and *gpmA* were only present in *Bifidobacterium* and *Lactobacillus* genomes. Some acid resistance genes such as *gpmL*, *bshA*, *bshB*, and *bshC* were only present in *Geobacillus* genomes. 8 bile resistance genes such as *nagB*, *pyrG*, *argS*, *rpsC*, *rpsE*, *rplD*, *rplE*, and *rplF* were present in all the genomes. Osmotic stress-related genes were absent in *Bifidobacterium* and *Lactobacillus* genomes, however, *opuD* and *opuC* were present in some *Geobacillus* genomes, and *opuCA*, *opuCB*, and *opuCC* were only present in *Geobacillus* genomes. Molecular chaperones such as *dnaK*, *dnaJ*, and *dnaG* were present in all the genomes whereas *groES* and *groEL* were present in some *Geobacillus* genomes only. 18 genes related to oxidative stress were screened and among them *taxA*. *TaxB*, *feoB*, and *msrC* were present in all the genomes whereas *ndhH*, *ndhB*, and *ndhC* were present only in *Geobacillus* genomes and *msrB* only in *Lactobacillus*. 7 genes related to vitamin biosynthesis were screened and genes *BtuD*, *copA*, and *copZ* were present in all the genomes whereas *BtuF* and *CsoR* were only present in *Geobacillus* genomes. Principle component analysis was done based on all the parameters (genes) related to probiotics including antibiotic resistance genes. The Principle component 1 (PC1) significantly shows the positive correlation between *Bifidobacteria* and *Lactobacillus* genomes and their relatively less positive correlation with the genomes of *Geobacillus* as shown in **Fig.6**.

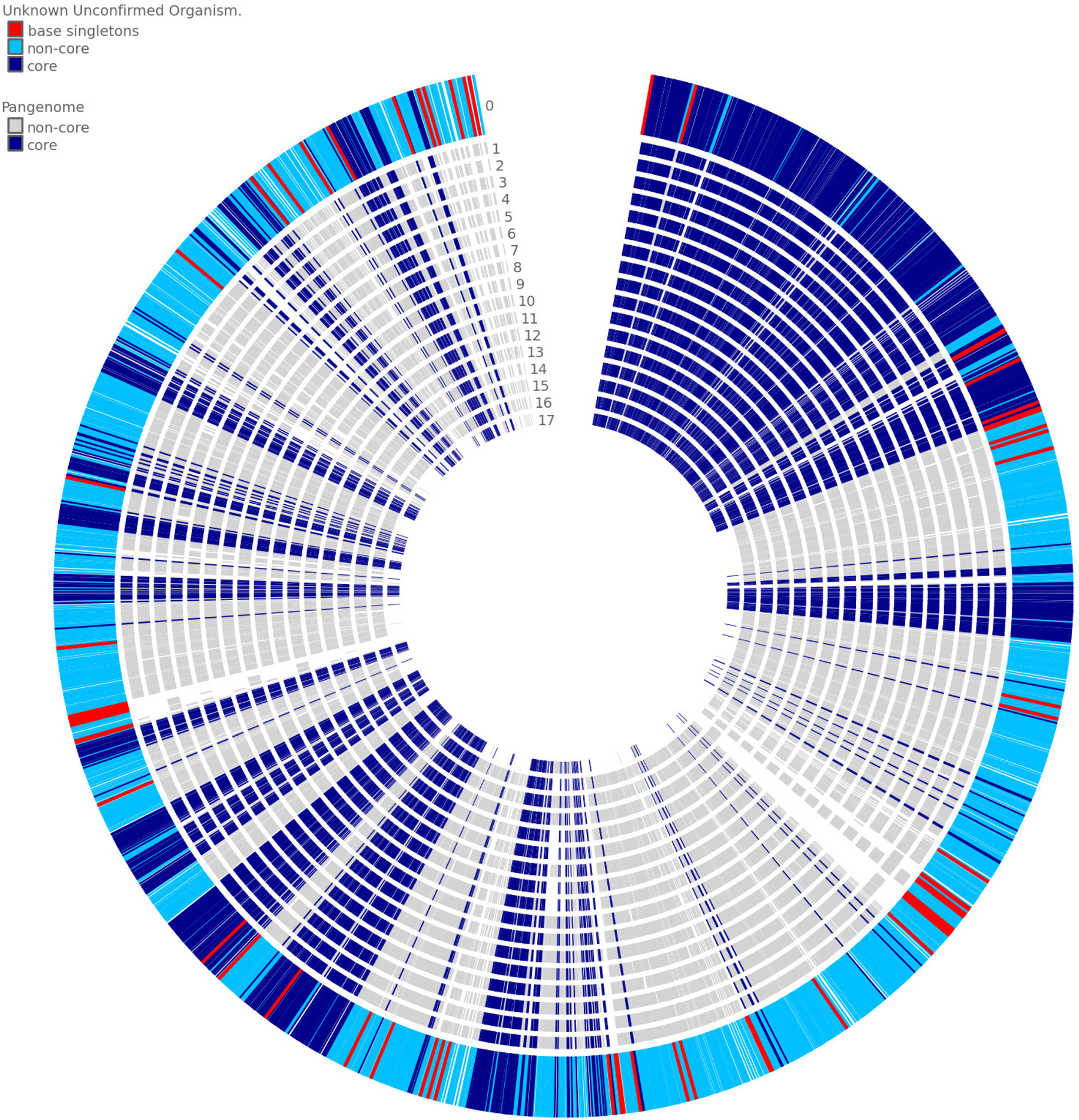

**Table 2.**
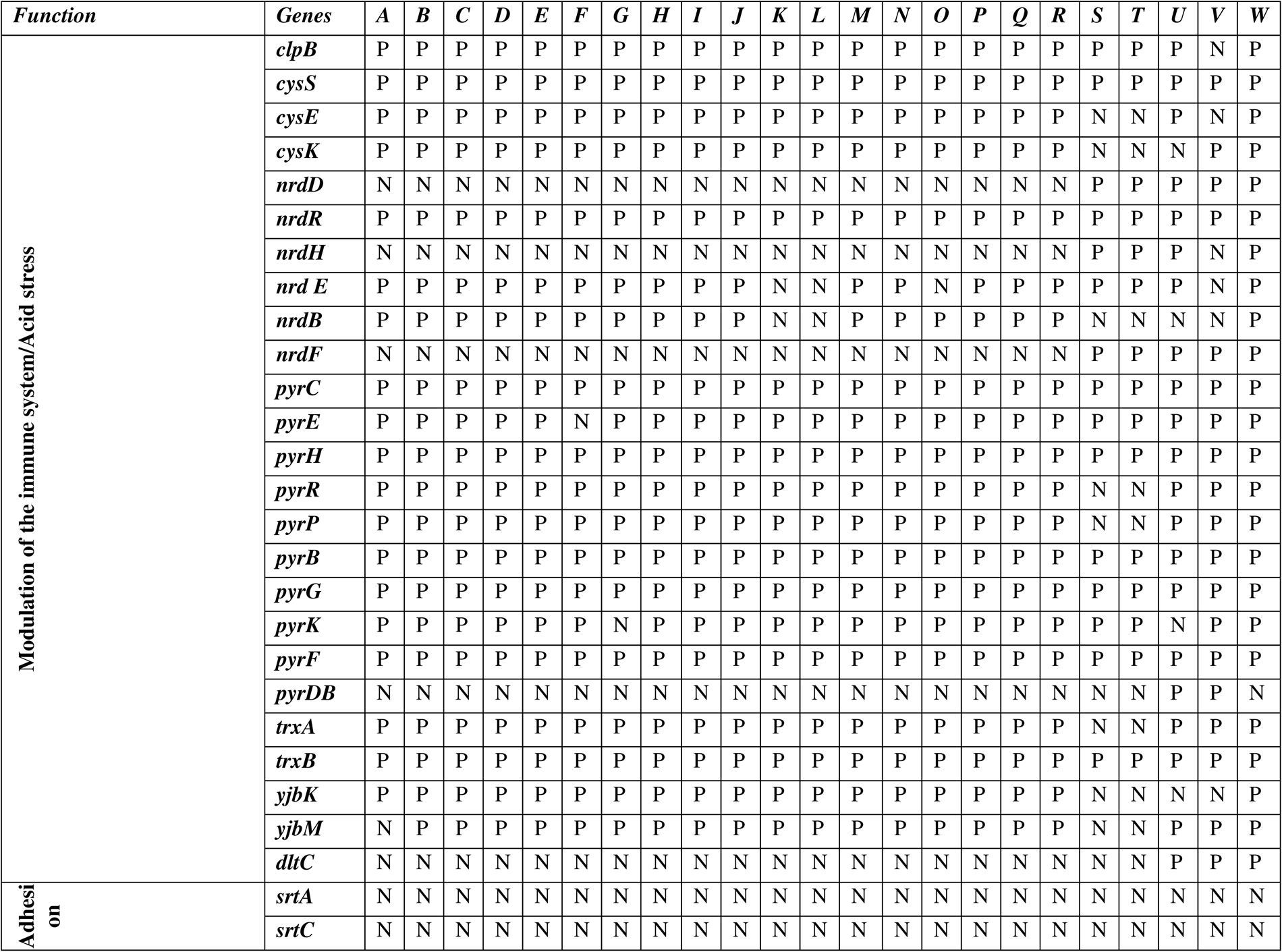
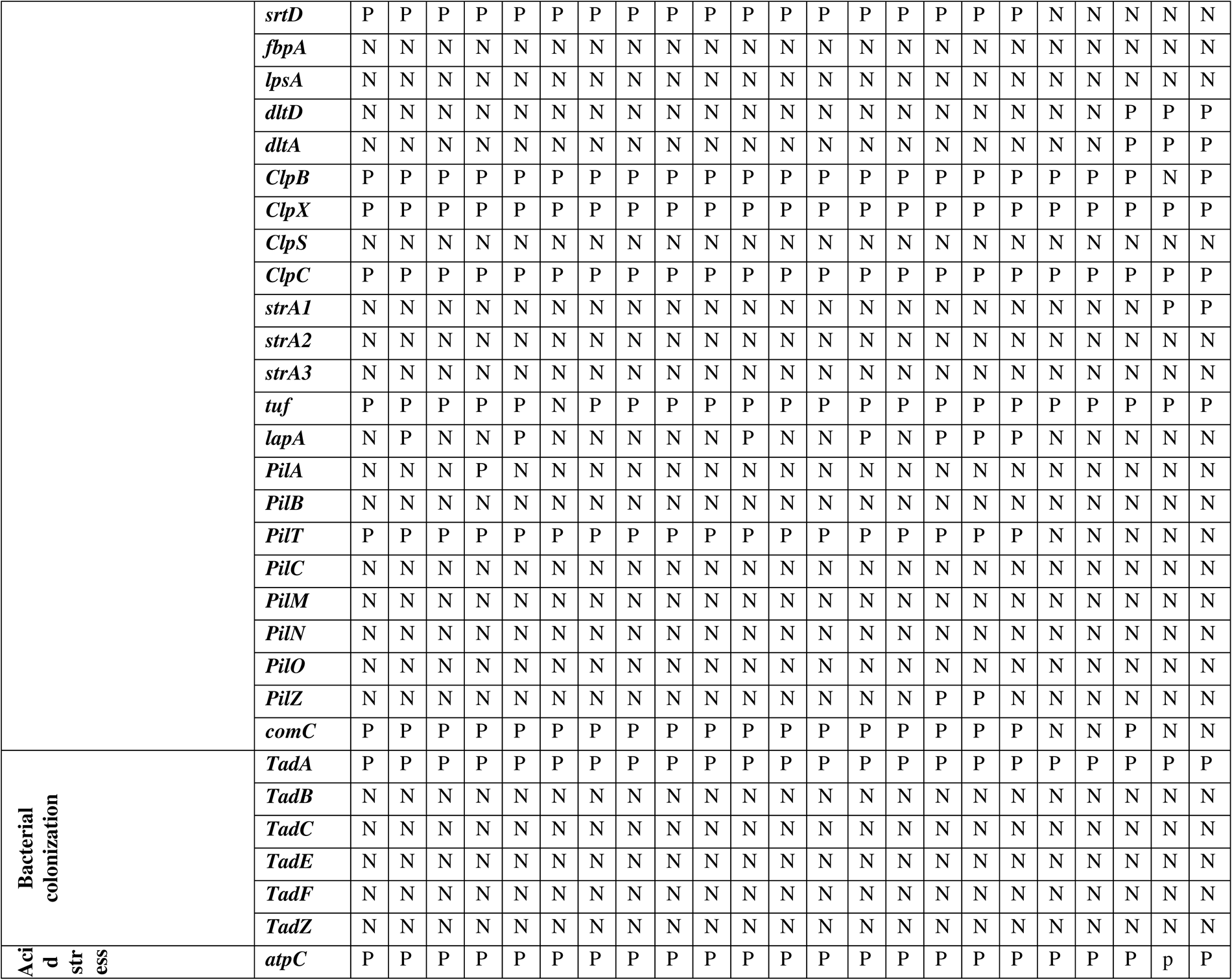
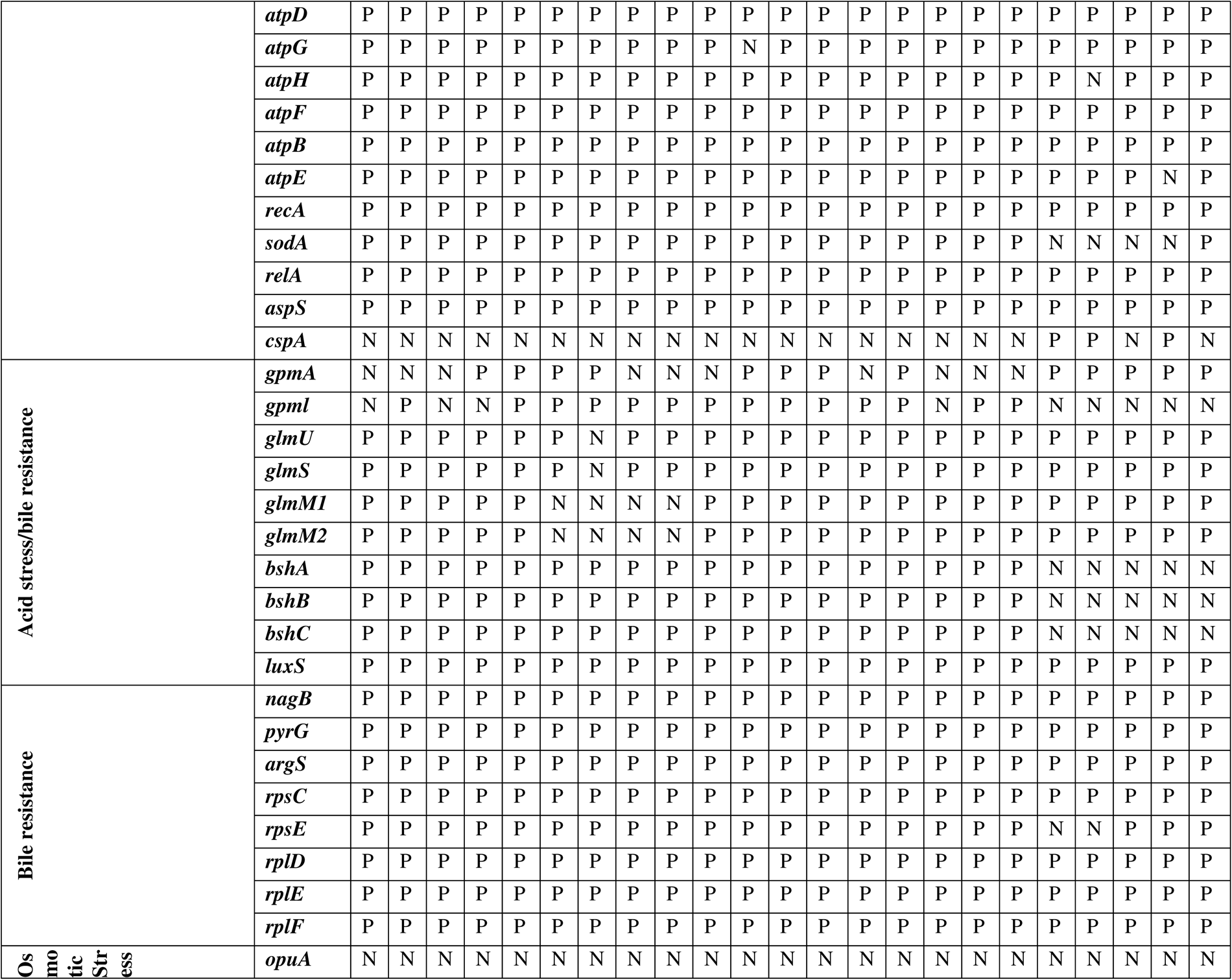
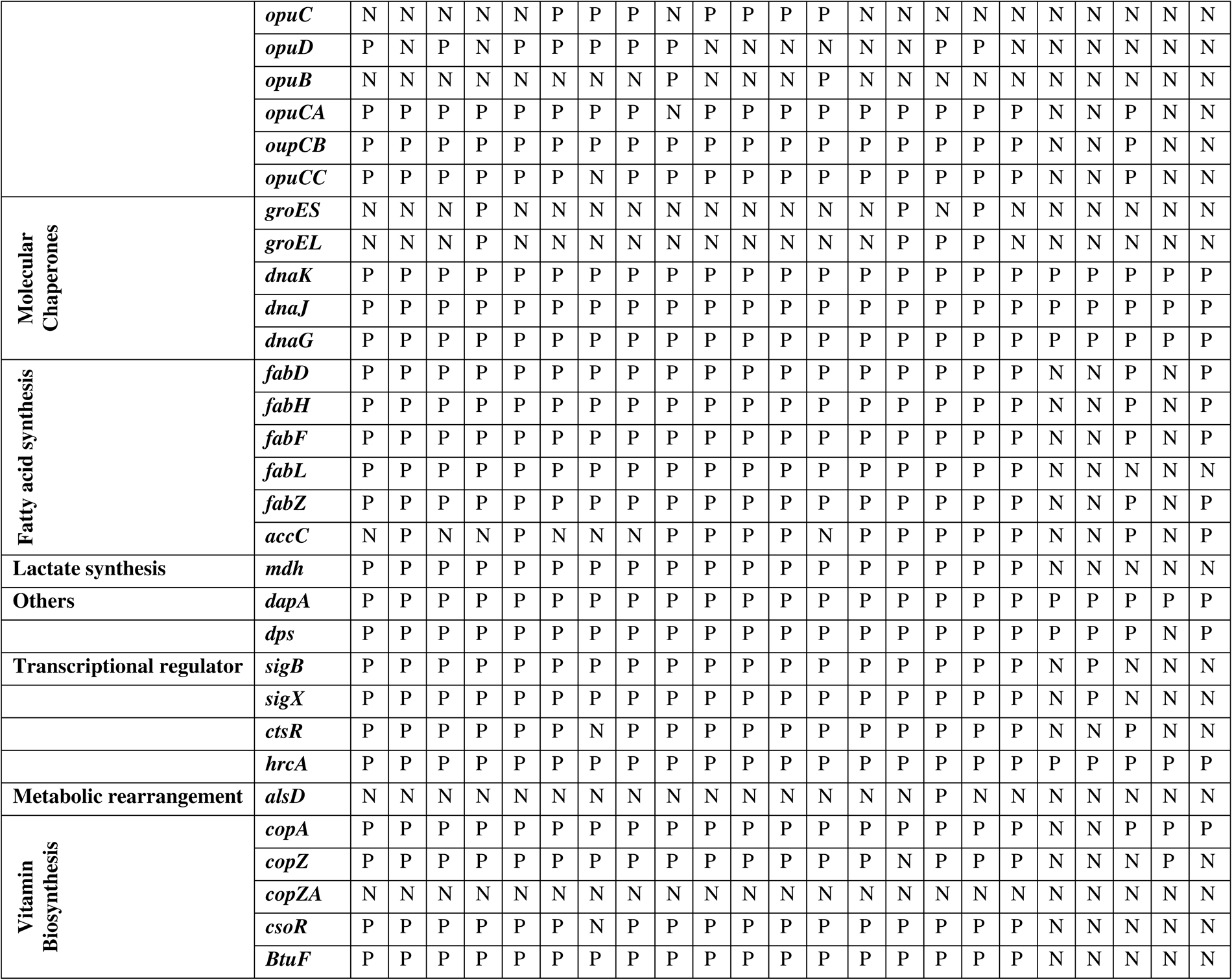
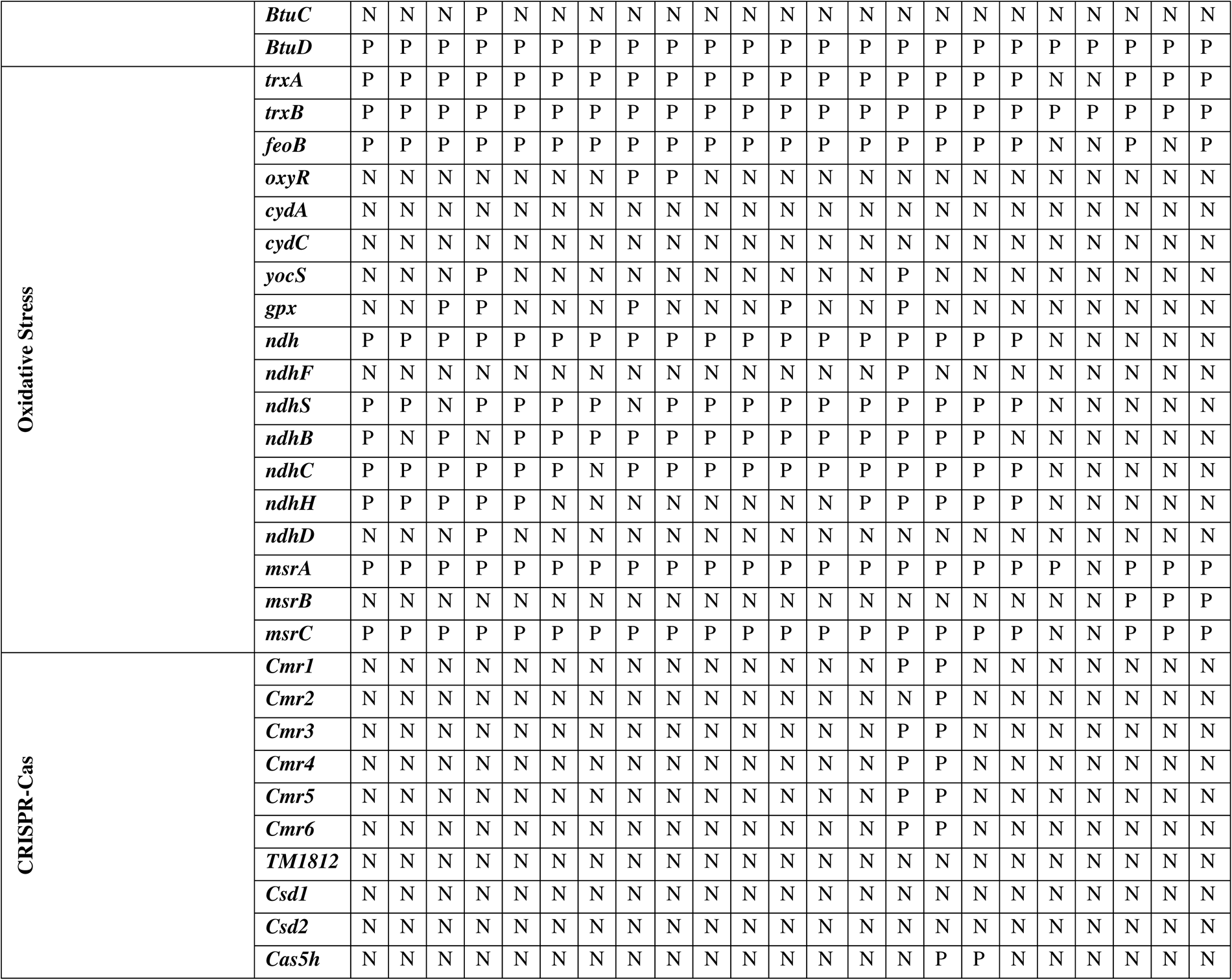
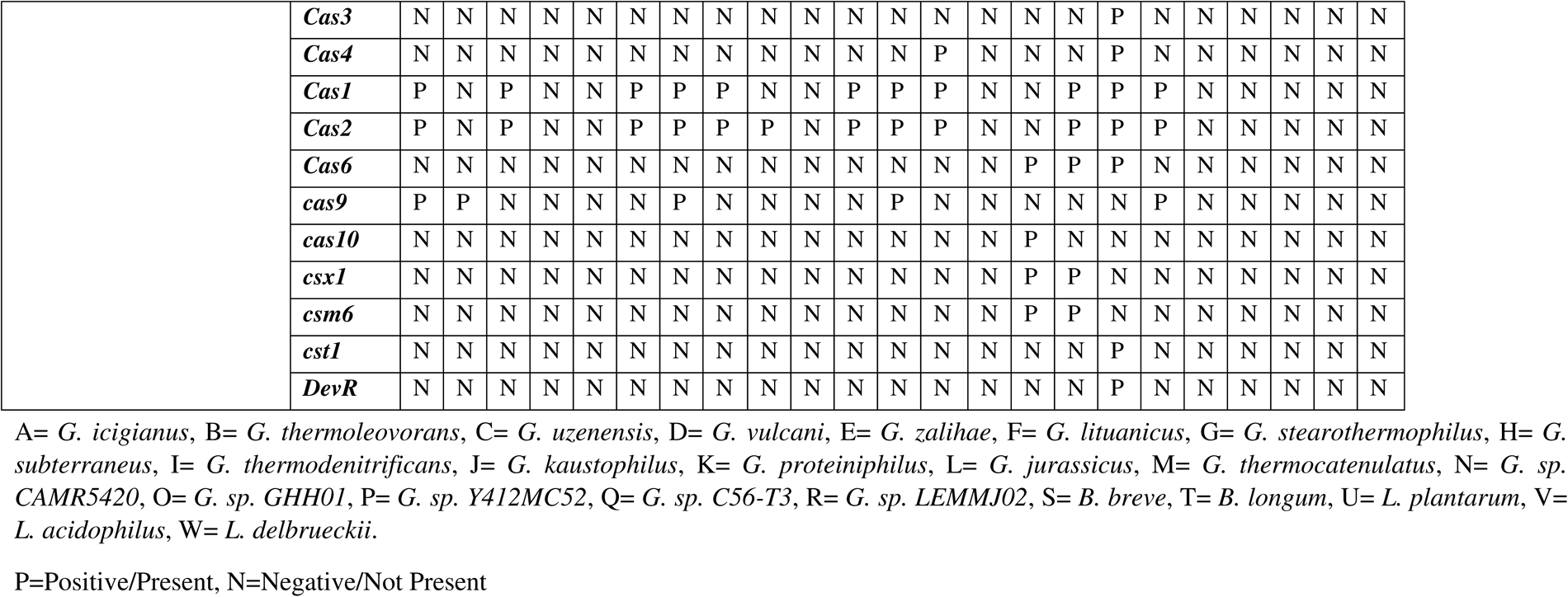
Prediction of genes related to probiotics.

### Assessment of Antibiotic Resistance Genes, Virulence Factors, and Toxins

The presence of antibiotic resistance genes was anticipated within the genomes under investigation. It was shown that only a few antibiotic-resistance genes such as *vanY*, *vanT*, and *rpoB* were present in all the genomes. However, other genes like *ImrD*, *rpsL*, and *qacJ* were present in *Lactobacillus* genomes as shown in **Table 3**. None of the toxin-related genes were present in *Geobacillus*, *Lactobacillus,* and *Bifidobacterium* genomes. Only a few toxin-like proteins such as *PhoH*, *MazF*, *HicA*, and *PemK* genes were predicted in some *Geobacillus* genomes. However, many genes coding toxins such as *FitB*, *PemK*, *MazF*, *ParB*, *RelB*, *RelE*, *MraZ*, *YoeB*, *PhoH*, and *HipA* toxin-related genes were present in genus *Bifidobacterium* and *Lactobacillus* species as shown in Table. 4. No virulence factors were found in any genomes studied as shown in Table. 3.

**Table 3.**
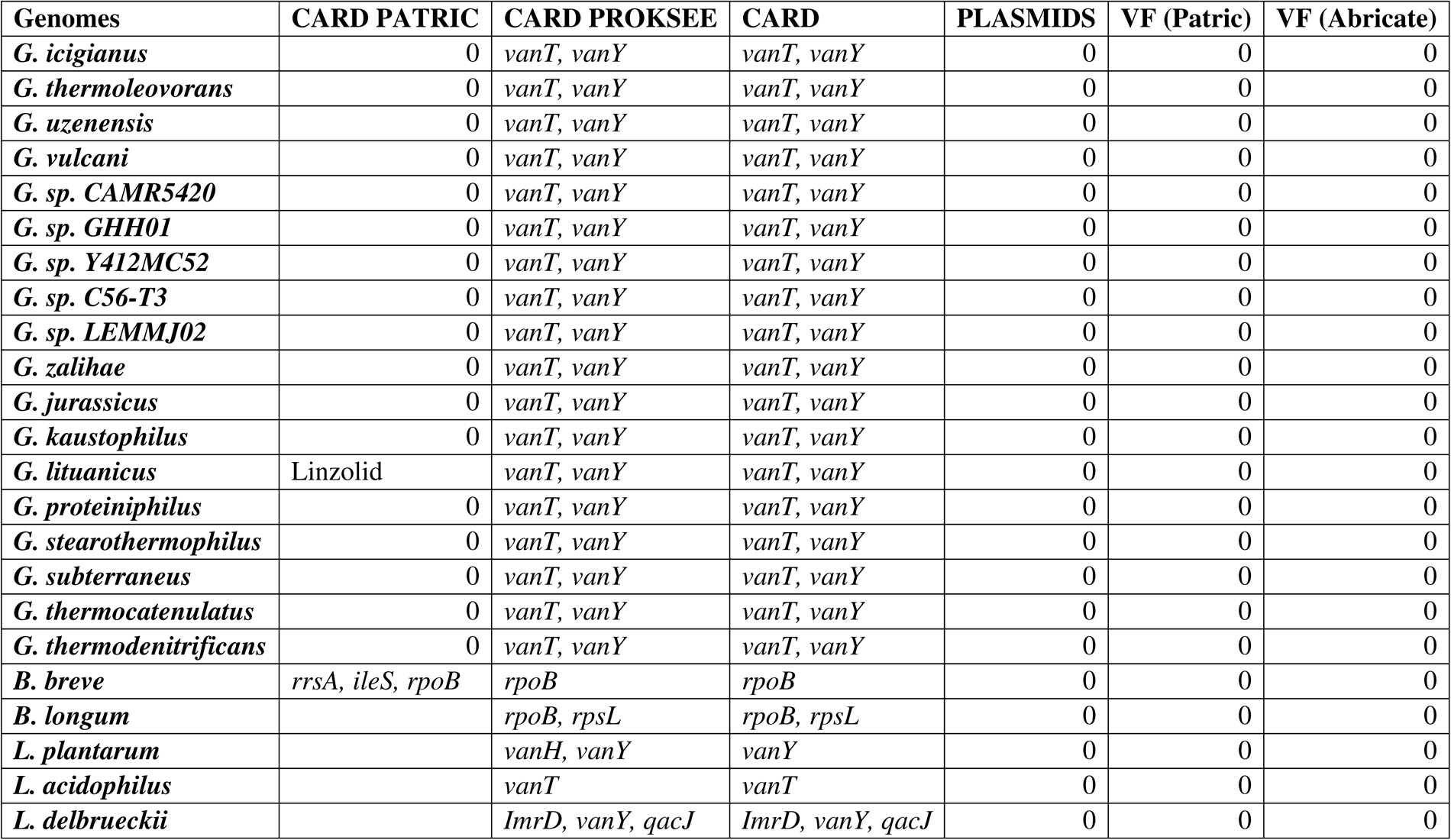
ARGs, Plasmids and Virulence factors present in studied genomes.

**Table 4.**
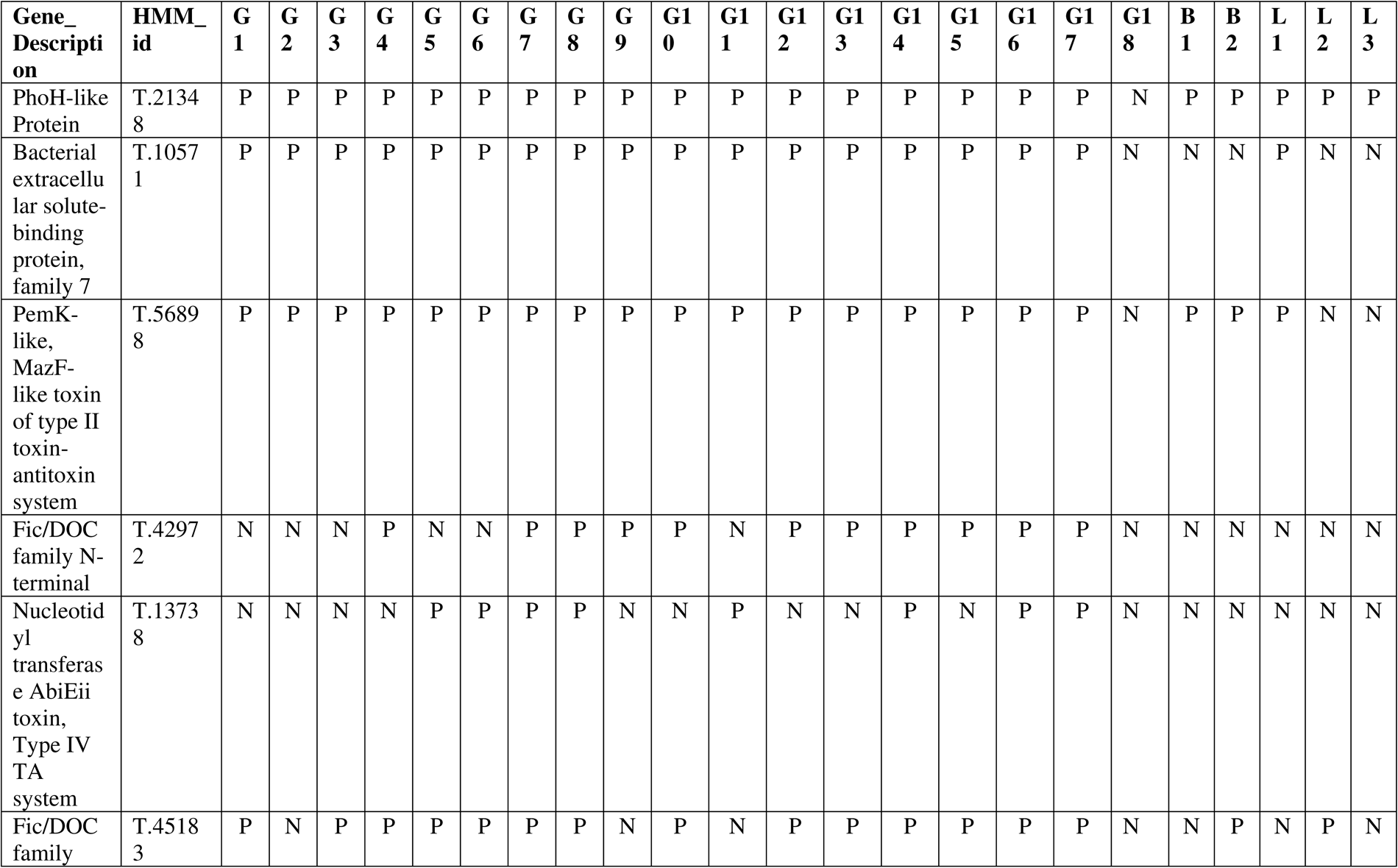
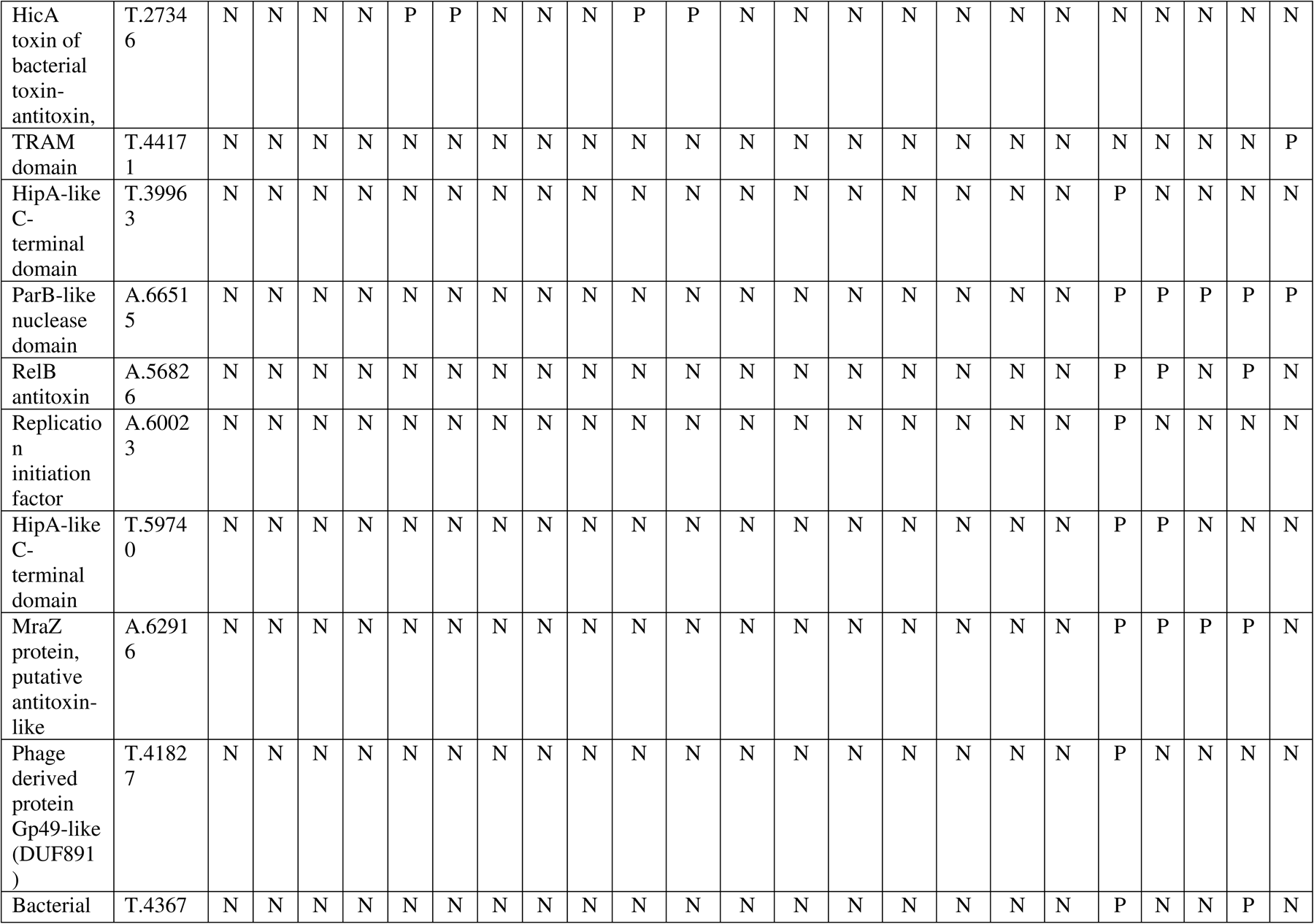
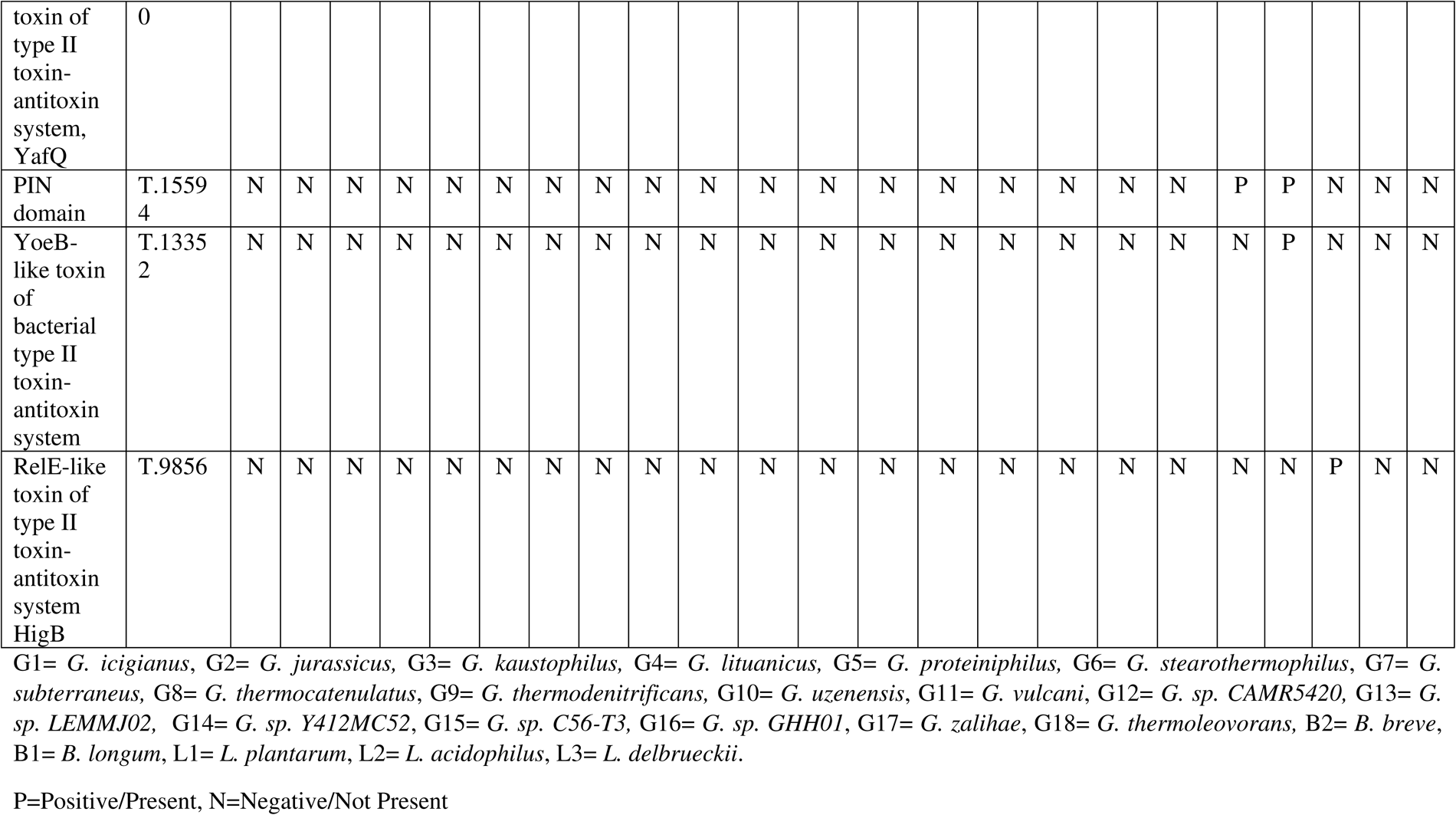
Presence of genes related to Toxins.

### Assessment of Mobile Genetic Elements, Insertion Sequences, Plasmids, Prophages, Bacteriocins, and CRISPR-Cas Systems

Mobile genetic elements were found to be absent in all genomes except a few mobile elements present in *B. brevis* such as IS/Tn. Putative insertion sequences were present in many *Geobacillus*, *Lactobacillus*, and *Bifidobacterium* genomes such as Putative IME, Putative ICE with T4SS, and Putative IME without identified DR as shown in **Supplementary Table. 1.** Plasmids were found to be absent in all the genomes Table. 3. Many bacteriocins such as Circularin_A, ComX1, ComX4, Salivaricin_D, Sactipeptides, Pumilarin, and Geobacillin_I_like were present in various genomes of *Geobacillus* like *G. icigianus*, *G. kaustophilus*, *G. lituanicus*, *G. C56T3*, *G. stearothermophilus*, *G. thermocatenulatus*, *G. vulcani,* etc. However, among the two *Bifidobacterium* genomes, only *B. longum* possesses a few bacteriocins such as Geobacillin_I_like and Propionicin_SM1. Moreover, *Lactobacillus* genomes possess different bacteriocins than that of *Geobacillus* and *Bifidobacterium* genomes such as Plantaricin_E/F/A/N/J/K, Enterocin_X, Acidocin_J1132, Enterolysin_A, and Bacteriocin_helveticin_J as shown in Table.5. Among 18 *Geobacillus* genomes, 11 possess prophages mainly within the family of Myoviridae and Siphoviridae. Similarly, *Bifidobacterium* and *Lactobacillus* possess prophages mainly from the family of Siphoviridae shown in Table. 6. CRISPR-Cas Systems were found to be present only in *Geobacillus* genomes. *Cas1*, *Cas2*, *Cas3*, *Cas4*, *Cas5h*, *Cas6*, *Cas9*, *Cmr1*, *Cmr3*, *Cmr4*, *Cmr5*, and *Cmr6* are present in many *Geobacillus* genomes such as *G. icigianus*, *G. thermoleovorans*, *G. uzenensis*, *G. stearothermophilus*, *G. subterraneus*, and *G. jurassicus* etc. However, CRISPR-Cas Systems were found to be absent in *Bifidobacterium* and *Lactobacillus* genomes as shown in Table. 2.

**Table 5.**
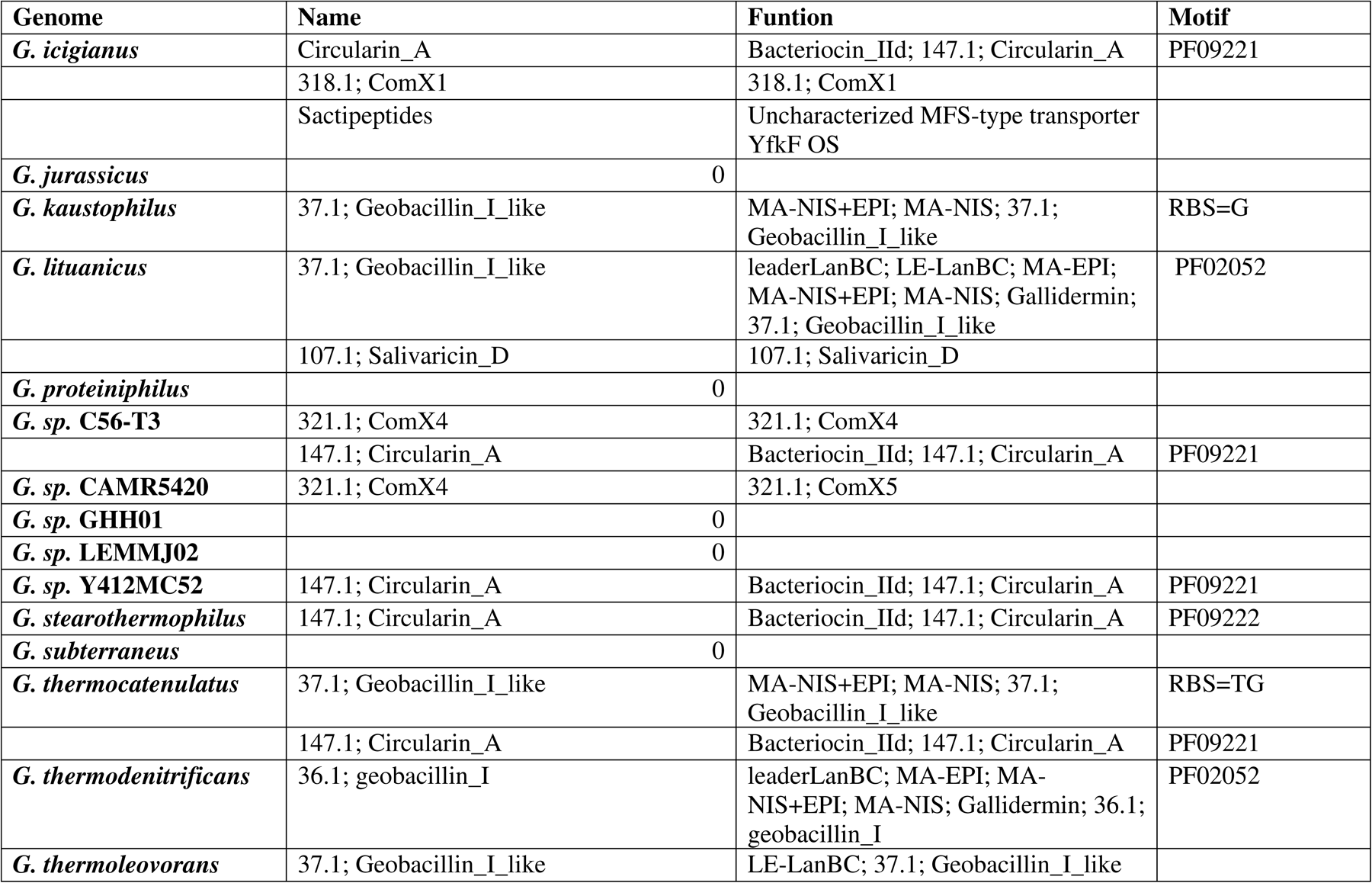
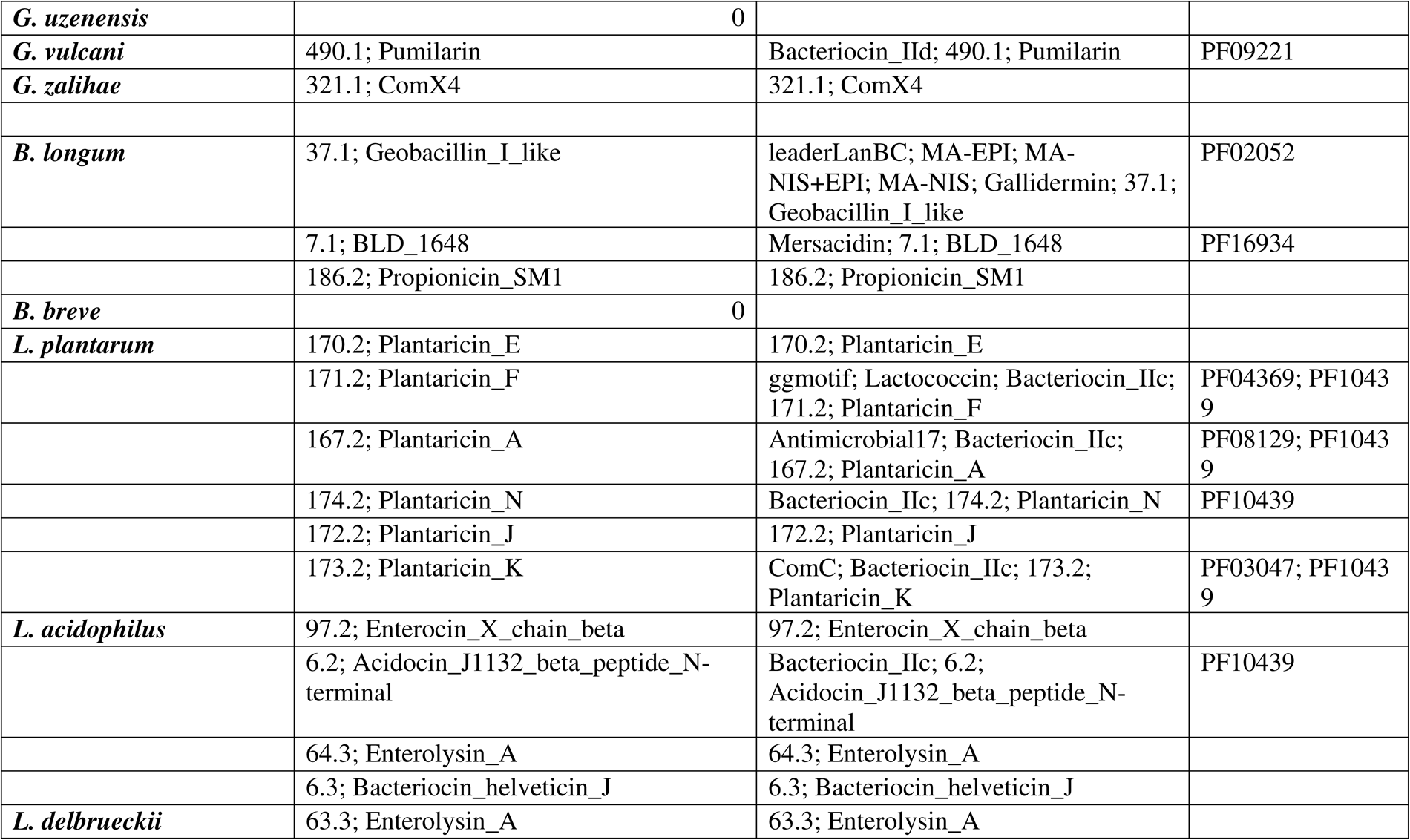
Presence of bacteriocins in studied genomes.

**Table 6.**
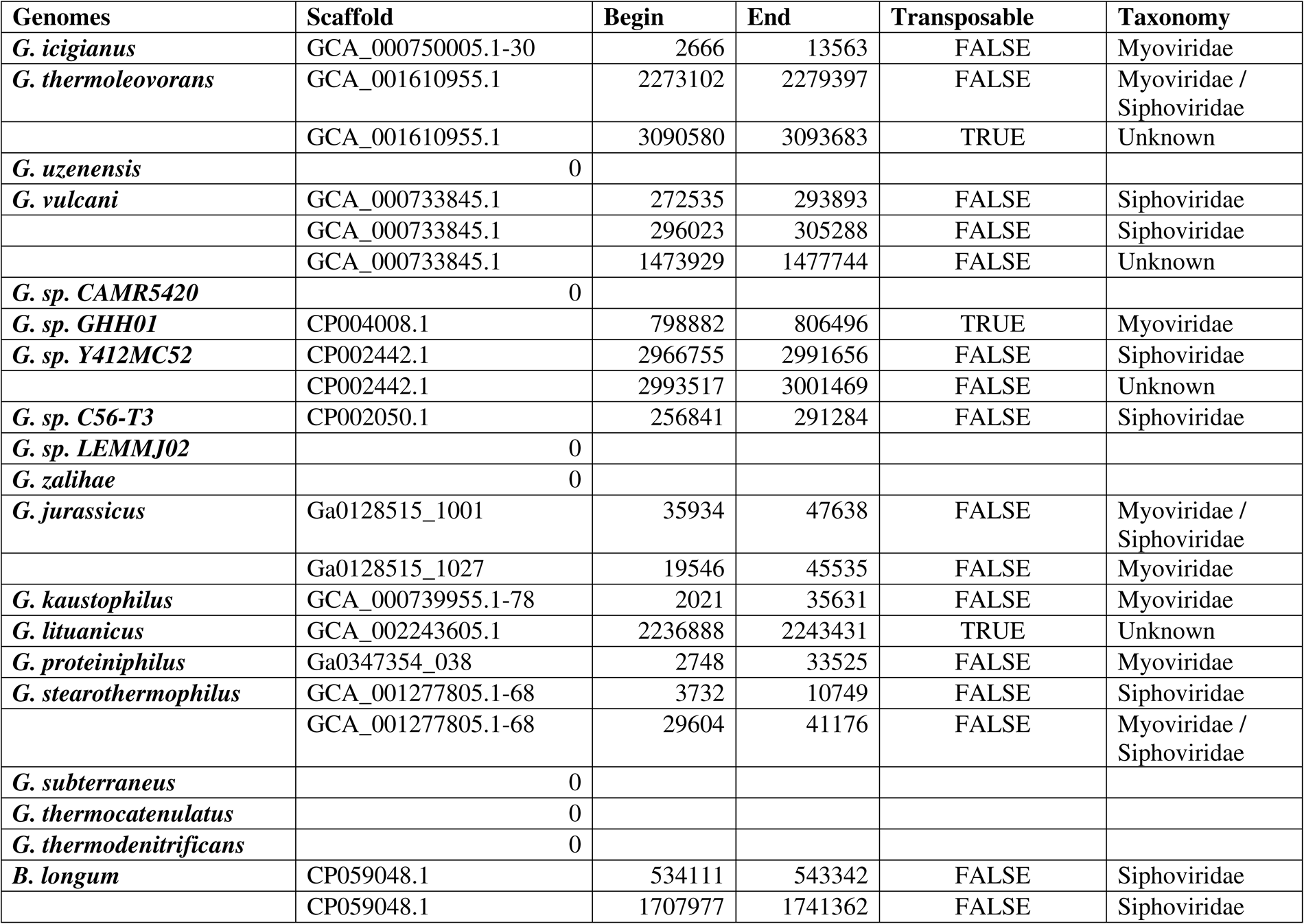
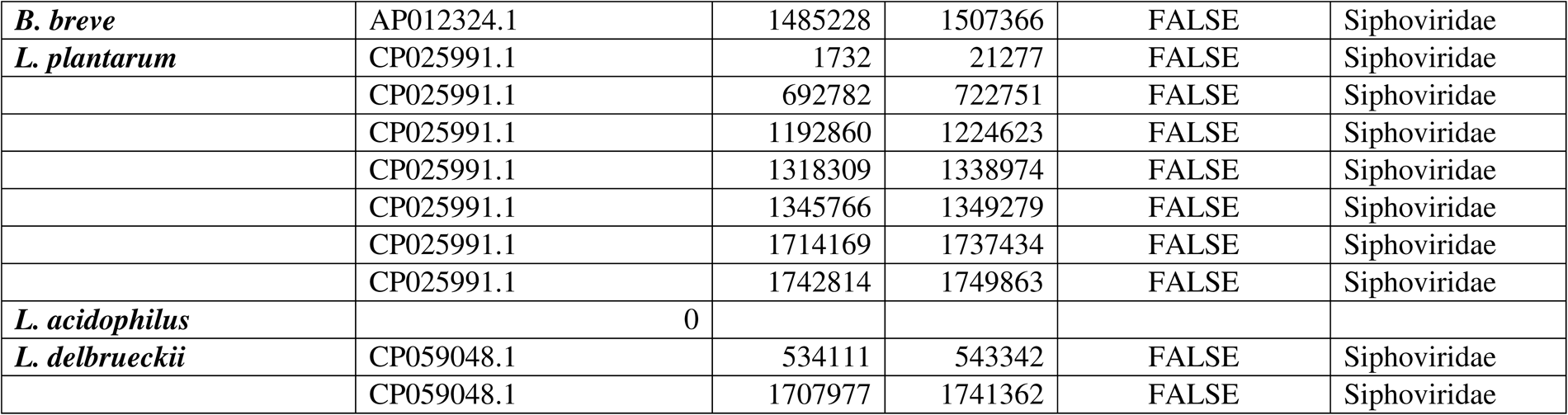
Presence of Prophages in studied genomes.

### Identification of Carbohydrate-active Enzymes (CAZyme)

Carbohydrate-active Enzymes (CAZyme) were found to be present in almost all genomes. Glycoside hydrolases were present in abundance and various glycoside hydrolase classes such as GH1, GH13, GH13_1, GH13_2, GH13_4, GH13_4, GH13_10, GH13_12, GH13_13, GH13_14, GH13_20, GH13_21, GH13_23, GH13_29, GH13_31, GH13_36, GH13_37, GH13_39, GH13_41, and GH18 were present with more than 2 hits or genes present. Among Glycosyl transferases GT2, GT4, GT51, and GT108 were present with more than 2 hits. CBM34 and CBM68 are present with more than 2 hits. CE4 and CE9 were the carbohydrate esterases present with more than 2 hits. Polysaccharide lyases were absent in all the genomes. AA1 and AA1_3 were CAZymes with Auxiliary activities having more than 2 hits. The above-mentioned CAZymes (except Polysaccharide lyases) were present in all the genomes as shown in Heatmap **Fig.7**.

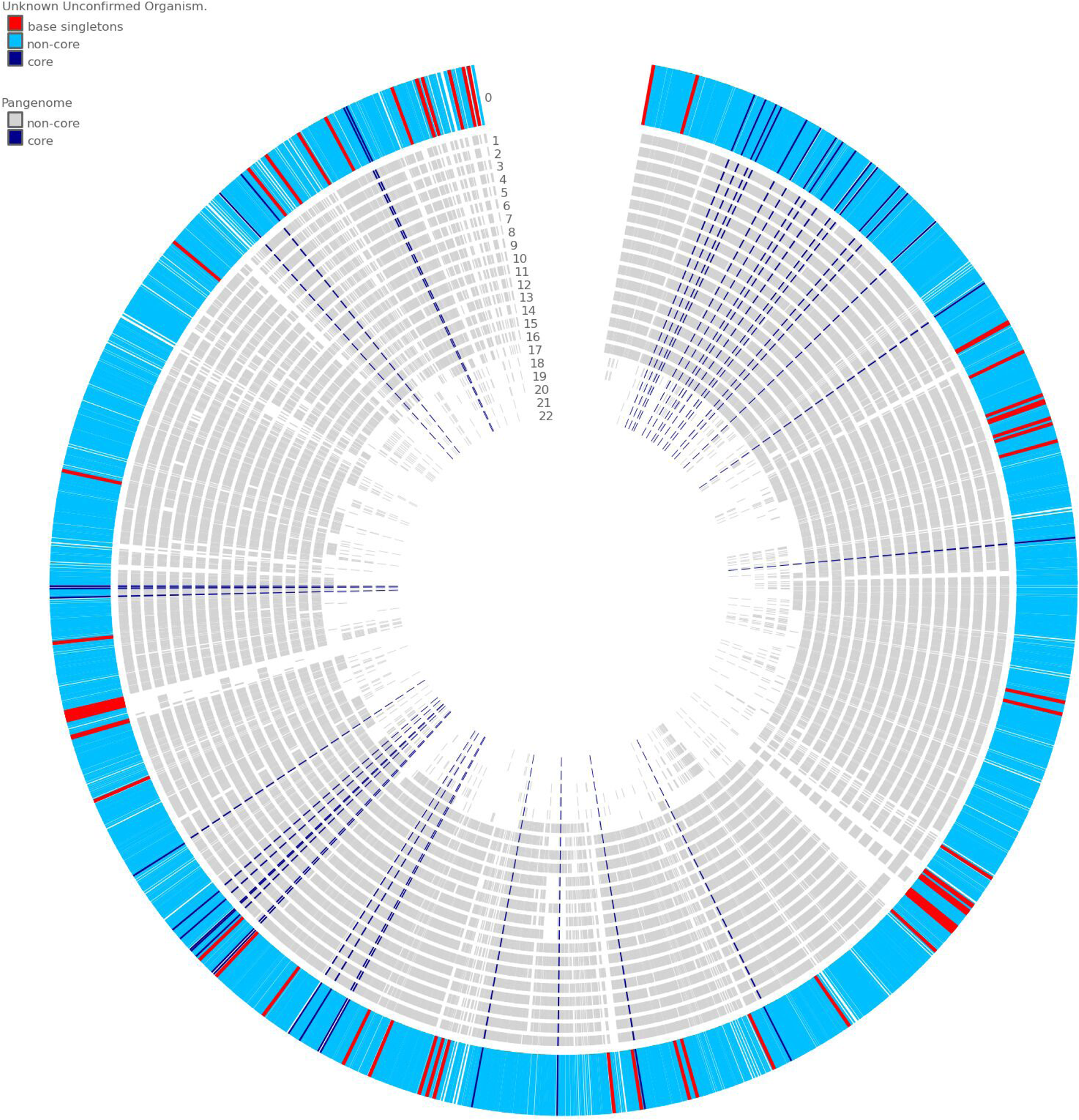

### Pangenome Analysi2s

The total number of genomes studied in the pangenome was 23 and among them 18 were *Geobacillus* species, 3 were *Lactobacillus* species and 2 were *Bifidobacterium* species. The pan-genome analysis was done within the genus *Geobacillus* alone and with other genera like *Lactobacillus* and *Bifidobacterium*. The pan-genome analysis of all 23 genomes represented 38393 protein-coding genes (with translation), 30702 genes in homolog families, and 7691 genes in singleton families. The total number of families was 11637, 3764 were homolog families, and 7873 were singleton families. When taking the pan-genome analysis of only the genus *Geobacillus* into consideration, the pan-genome represented 25284 protein-coding genes (with translation), 23504 genes in homolog families, and 1680 genes in singleton families. The total number of families was 4641, 2813 were homolog families and 1828 were singleton families **Supplementary Table 2**. The core pangenome rarefaction curve of the genus *Geobacillus* shows the increasing number of gene clusters with every added genome. Thus, this shows that the *Geobacillus* pan-genome is an open pangenome as shown in **Fig.8**. The pangenome circle plot was contrived to plot the core, non-core, and singletons. The circle plot of the genus *Geobacillus* shows larger core genes than that of singletons shared between the genomes whereas when the circle plot was made between *Geobacillus*, *Lactobacillus*, and *Bifidobacterium* genomes, they shared very few core genes as shown in **Fig.9; Fig 10, Supplementary Fig. 2; Fig 3**. Phylogenetic Pangenome Accumulation was run to view the pangenome in a phylogenetic context and to determine the entry and exit of gene families in the branch of interest. The Phylogenetic Pangenome Accumulation tree shows the bifurcation of two main clades as shown in the phylogenetic trees discussed above. Node zero was important as it bifurcates into two main clades with node 1 (gives rise to genus *Bifidobacterium* clade) and node 6. Node 6 was also important as it bifurcates into two separate branches of the genus *Geobacillus* as one clade and the genus *Lactobacillus* as another clade. At node zero, there was the maximum number of total genes present (17392) with a very less perfect core of only 0.8%. there is the accumulation of genes at node 1 for genus Bifidobacterium with an increase in perfect core of 27.3%. Similar to node zero there was the second maximum number of total genes (14861) (as the rest of the genes got accumulated into genus *Bifidobacterium*) with a very small perfect core of only 1.6%. This total gene numbers gradually accumulated and shows the increase in perfect core as shown at node 18 (with the perfect core of 8.6%) for the genus *Geobacillus* and at node 7 (with a perfect core of 8.9%) for the genus *Lactobacillus.* These genes further get accumulated within sub-branches as shown in **Fig.11**.

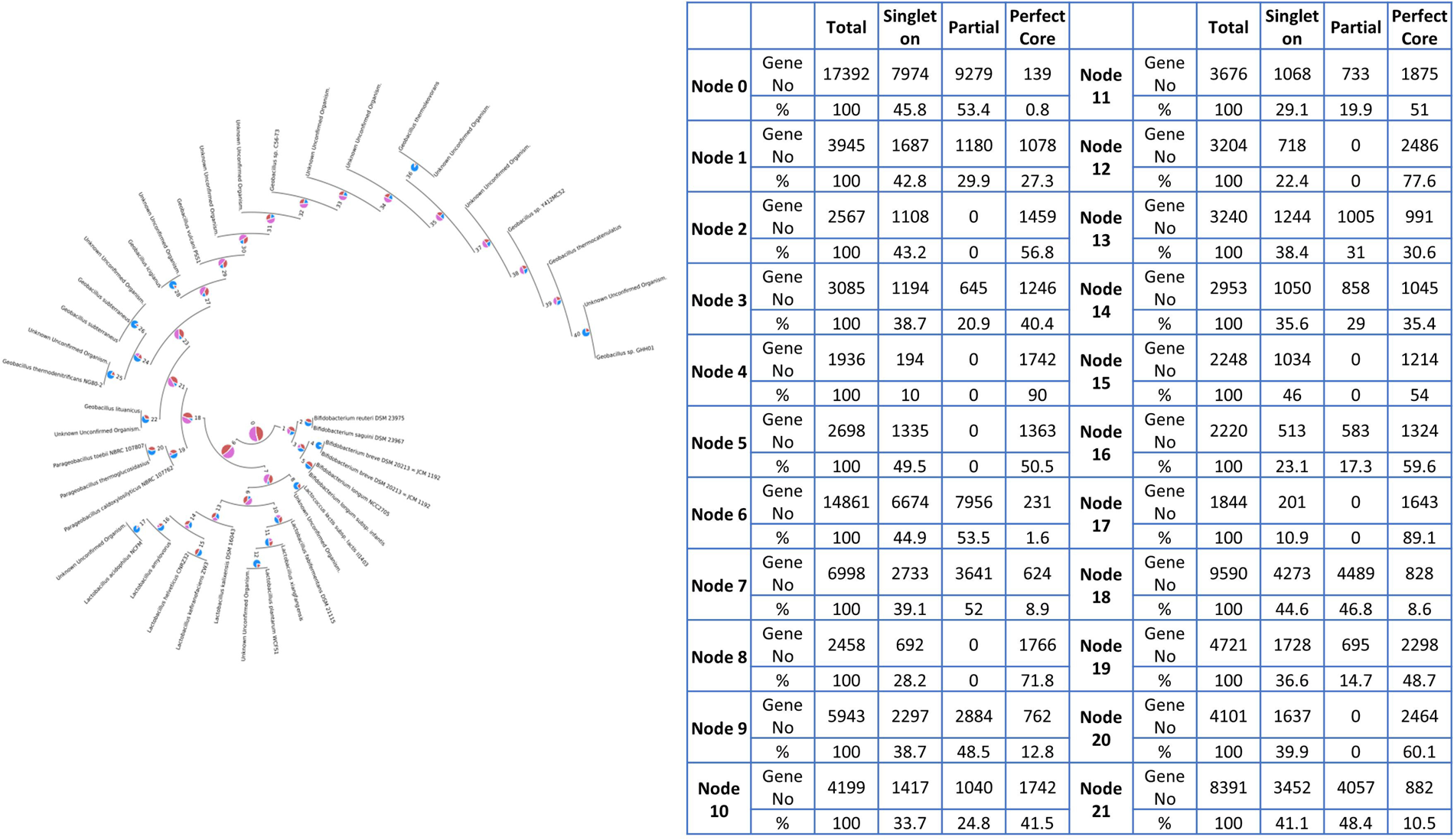

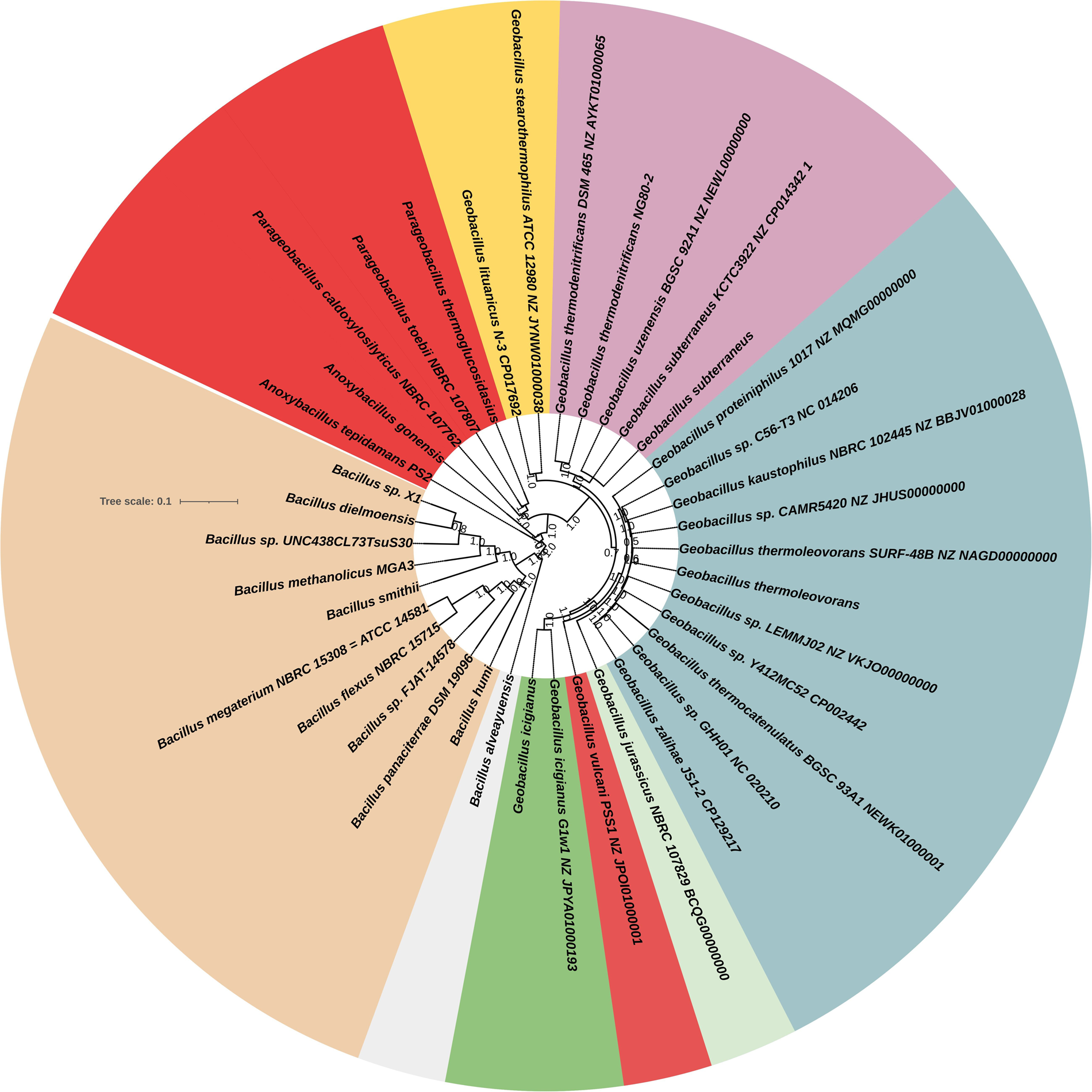

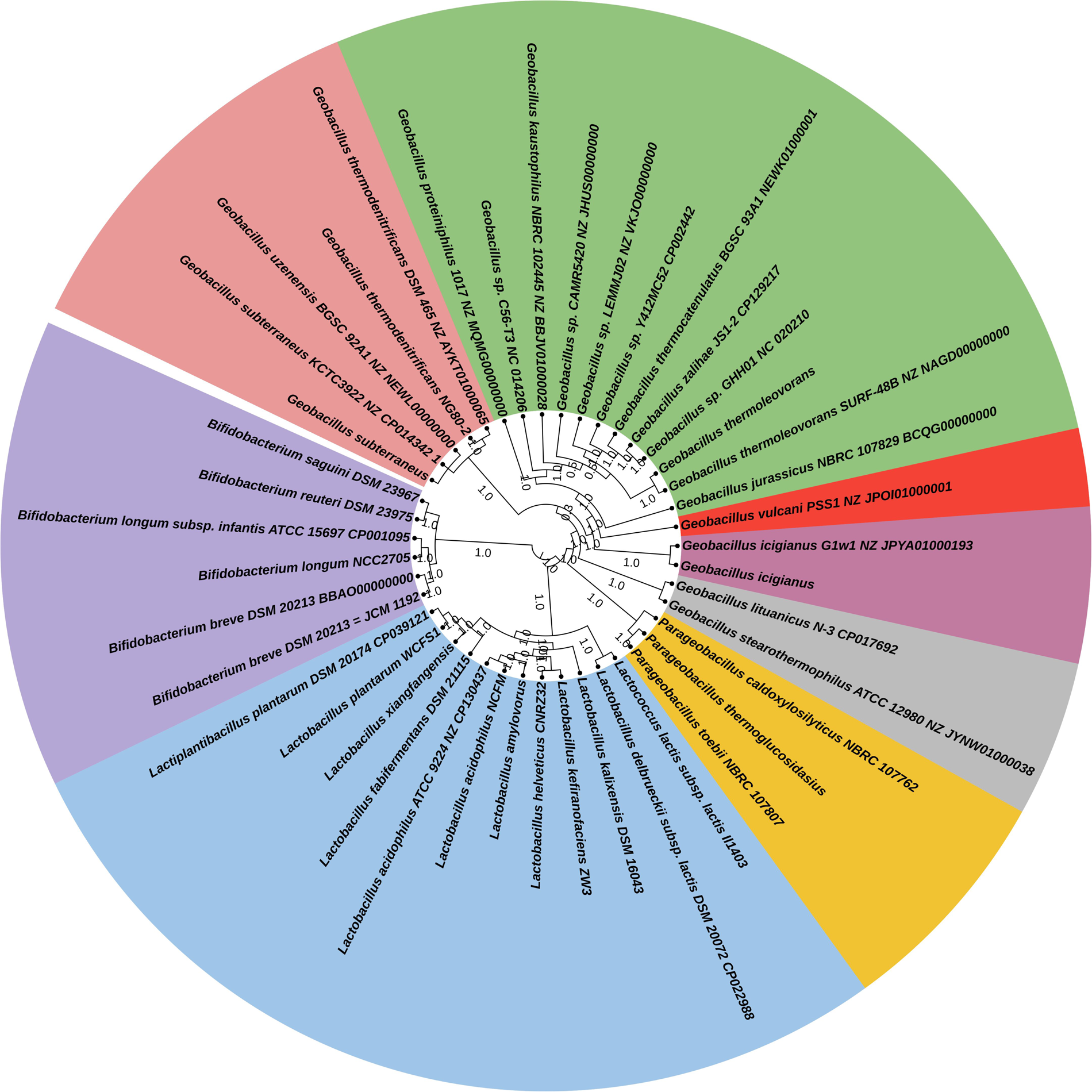

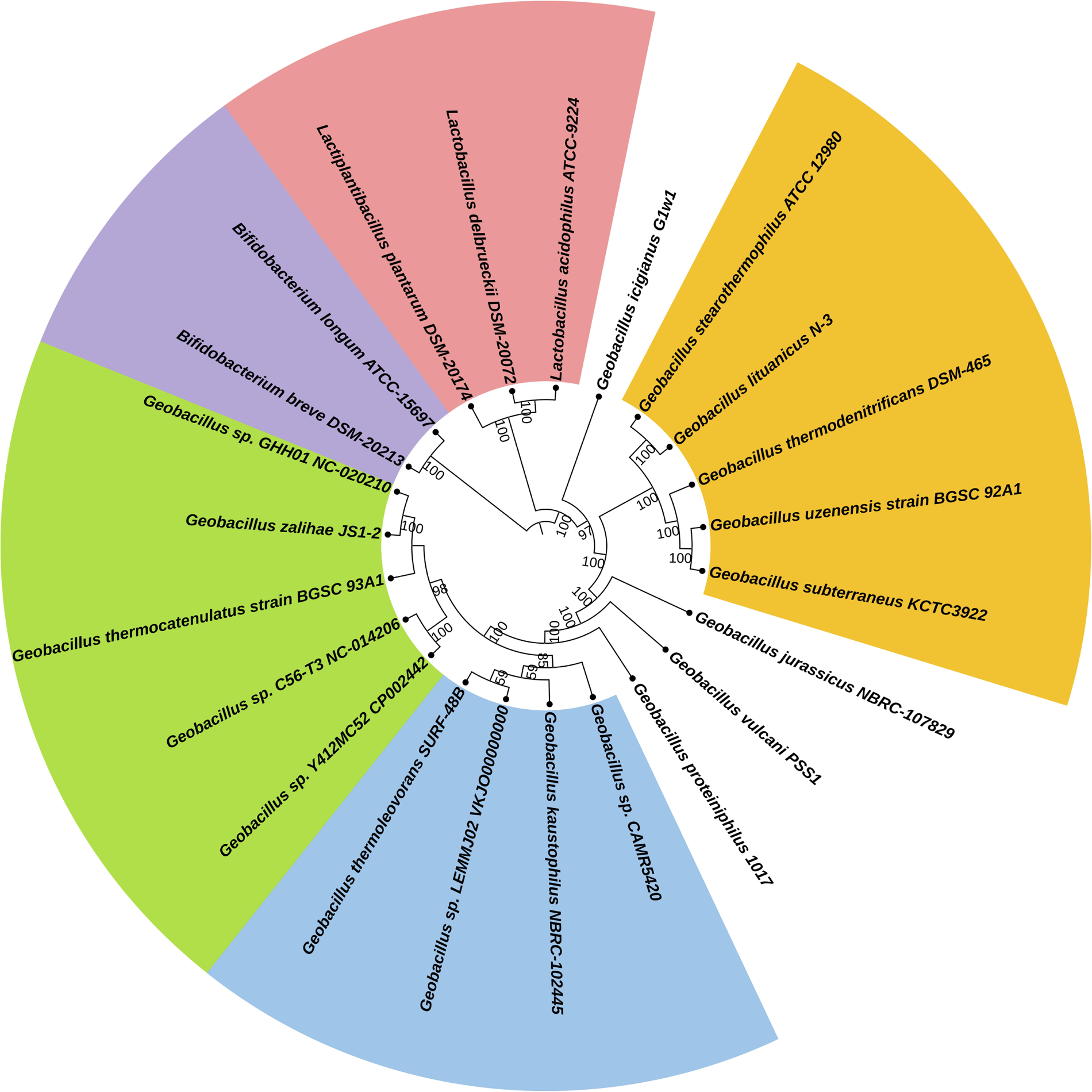

## Discussion

The species of the *Geobacillus* genus exhibits ecological, physiological, and genetic diversity, merely due to micro-evolutionary traits such as horizontal gene transfer, which remains a strong factor contributing to the evolution of *Bacillus* to *Geobacillus* (26) Thriving in various harsh conditions thermophiles have been exploited in various industrial and biotechnological applications. However, only a few studies suggest the probiotic nature of the species belonging to the genus *Geobacillus*. Here we have presented the vast *in silico* evaluation of the genus *Geobacillus* considering its probiotic nature. The analysis has been done in genus *Geobacillus* keeping already known probiotic strains of genus *Lactobacillus* and *Bifidobacterium* as positive controls. Various characteristics studied so far for claiming any probiotic species have been covered in this study.

Taxonomic and phylogenetic studies revealed that the genomes of *Geobacillus* possess high nucleotide identity and thus are correctly represented in the genus *Geobacillus*. Similar phylogenetic results have been shown earlier for genus *Geobacillus* (27). When considering two other genera such as *Lactobacillus* and *Bifidobacterium*, the phylogenetic trees based on COG-based core genes and single-copy genes using single-copy PGFams specifically form separate clades of each genus thus confirm their correct representation within their respective genus. The annotation results using Prokka suggested that the subsystems highly represented were replication, recombination, repair; post-translational modifications, protein metabolism, and chaperones; nucleotide transport and metabolism; signal transduction mechanism; general functions; energy production and conversion; translation, ribosome structure, and biogenesis; carbohydrate metabolism, amino acid metabolism, and transcription. Whereas the unrepresented features were secondary metabolites biosynthesis, transport, catabolism, intracellular trafficking, secretion, chromatin structure and dynamics, cytoskeleton, and RNA processing and modification. Our research demonstrates that the core genome encompasses the majority of the COG categories. However, flexible genome (accessory and singleton) also shows a considerable percentage as shown in pan-genome accumulation results. Consequently, comprehensive comparative analyses unveiled that crucial functional classes and essential housekeeping genes remained constant within the core genome. Conversely, genes associated with environmental interaction or the synthesis of secondary metabolites notably exhibited higher abundance within the pan-genome. Also, the considerable flexible genome percentage reveals that Horizontal Gene Transfer (HGT) could represent a crucial mechanism contributing to the environmental adaptive characteristics observed in the genus *Geobacillus*.

Similar findings were reported by Bezuidt et al. in 2016, indicating an overabundance of genes related to COG categories including translation, ribosomal structure, coenzyme transport and metabolism, nucleotide transport and metabolism, as well as protein turnover and chaperones within the conserved core. (26). In a study conducted by Wang et al. in 2020, detailed comparative analyses indicated that core genome stability persisted in fundamental functional classes and essential housekeeping genes, while the flexible genome exhibited a higher prevalence of genes associated with environmental interaction or energy metabolism. Furthermore, instances of horizontal gene transfer (HGT) were identified among various *Geobacillus* species and thus the *Geobacillus* evolution appears to be influenced by environmental factors (28).

As in our study, we found that secondary metabolite metabolism is mainly confined to a flexible genome. Thus, to evaluate this metabolite potential is very important for industrial and biotechnological applications. Wang et al, 2020, shows that various genes for starch, arabinose, glucose, mannose, galactose, and xylose metabolism were also observed in the pan-genome, with a high occurrence of ABC-type sugar transporters that aid in sugar uptake. Therefore, the pan-genome analyses in *Geobacillus* reveal it as a good candidature for hemicellulose degradation (28). A similar overrepresentation of the genes was observed in a study conducted by (26) on pan-genome analyses of 29 *Geobacillus* genomes, depicting that such diverse metabolic machinery aids in its diverse biotechnological potential.

There are various measures for the assessment of any species to be probiotic. The FAO and WHO (2002) guidelines, alongside regulations from the Food and Drug Administration and the Ministry of Public Health in Thailand, delineate the criteria for utilizing probiotic microorganisms in food. These standards encompass specific criteria such as accurate identification, evaluation of probiotic characteristics (such as resistance to gastric acid and bile salts, adherence to mucosal surfaces and epithelial cells, and bile salt hydrolase activity), and a safety assessment encompassing factors like antimicrobial resistance (AMR), toxin production, and hemolytic activity (29, 30). We have evaluated and covered almost all the characteristics of genus *Geobacillus* to assess its probiotic potential concerning genus *Lactobacillus* and *Bifidobacterium*. Genes related to tolerance to various stress conditions, adhesion, colonization, immune response, etc. governing the probiotic potential were determined. Several genes associated with adhesion, such as *srtD*, *clpB*, and *clpC*, were identified, potentially playing a role in binding to cells or mucosal surfaces within the gut. (31, 32). Other genes such as *TadA*, *PilT*, *PilZ*, and *lapA* essential for pili structure, and thus bacterial colonization were found in the genus *Geobacillus* (33). The key feature defining a bacterium as a probiotic strain hinges on its ability to survive, adapt, or resist low-pH environments (34). Studies suggest that probiotic microorganisms harbor particular genes that aid in enduring or resisting adverse conditions (34). *Lactobacillus* strains typically exhibit resilience to acid and bile, allowing these bacteria to endure and adjust to harsh conditions, rendering them promising candidates for probiotic applications (35). Here we have detected various genes for acid and bile resistance in genus *Geobacillus* such as *sodA*, *luxS*, *glmU*, *atpH*, *recA*, etc. The F1F0-ATPase, encoded by the *atp* operon, typically comprises genes—namely, *atpB*, *atpE*, *atpF*, *atpH*, *atpA*, *atpG*, *atpD*, and *atpC*—in most microbes (36). While most of these genes were identified in the *Geobacillus* genome. These *atp* genes play a crucial role in enabling host microorganisms to survive or withstand acidic environments. The main role of the “*atp*” operon is to facilitate proton pumping, transporting protons from the bacterial cytoplasm outward, and contributing to the maintenance of a neutral pH in the bacterial cytosol. (37).

In response to hyperosmotic stress, organisms accumulate osmotically active compounds. These compounds, known as compatible solutes, do not hinder enzymatic processes within the cell. The gathering of these solutes, known as osmoadaptation, works to counter the outward movement of water, thereby maintaining cell turgor (38). Numerous thoroughly characterized compatible solute transporters have been detected in various gram-positive organisms, such as *Bacillus subtilis* and *Lactococcus lactis*. *OpuA*, *OpuB*, and *OpuC* belong to the ATP binding cassette (ABC) superfamily and are closely associated transporters. These transporters have the capability to transport distinct compounds: *OpuA* transports proline betaine and glycine betaine, *OpuB* transports choline, while *OpuC* transports ectoine, crotonobetaine, γ-butryobetaine, carnitine, choline-O-sulfate, choline, proline betaine, and glycine betaine. (39). These osmotic stress-related genes were absent in *Bifidobacterium* and *Lactobacillus* genomes, however, *opuD* and *opuC* were present in some *Geobacillus* genomes, and *opuCA*, *opuCB*, and *opuCC* were only present in *Geobacillus* genomes. Genes related to oxidative stress such as thioredoxin (*trx*), and ferrous iron transporter (*feoB*) were present in almost all the genomes. However, transcriptional regulator (*oxyR*), and NADH (*ndhH*, *ndhB*, and *ndhC*) were present only in *Geobacillus* genomes. These genes are associated with antioxidant and oxidative stress responses in many bacteria (40, 41).

The genes responsible for encoding chaperones (*dnaK*, *groEL*, *groES*) play a crucial role in a broad stress response, involving protection, elimination of damaged proteins, and various related functions. Additionally, the analysis revealed *dnaJ* and *grpE*, which have demonstrated a responsive behavior (upregulation) specifically to acidic environments (42). All the genomes examined contained these molecular chaperones. However, *groES* and *groEL* were present in some *Geobacillus* genomes only. The presence of these probiotic-related genes in the genus *Geobacillus* makes them a promising candidate for exploiting them as probiotic strains. However, culture-dependent analysis of these probiotic features must be validated.

Analyzing the mobilome offers a deeper comprehension of genome stability, adaptability, and evolution in host probiotics, particularly in evaluating the potential acquisition and transfer of new genes, including those linked to antibiotic resistance. Nevertheless, if a probiotic strain’s genome harbors mobile genetic elements (MGEs) along with antibiotic resistance genes, it becomes unfit for use due to the risk of transferring these resistance genes via processes like conjugation or other mechanisms (43, 44). The lack of mobile genetic elements, virulence factors, and minimal putative insertion sequences indicates the stability of the genus Geobacillus, potentially making it favorable for probiotic candidacy. Regarding the anticipated prophages found in *Geobacillus* genomes, only a small number belonging to the Myoviridae and Siphoviridae families were identified. However, most of these were hypothetical and lacked the typical genes responsible for encoding structural proteins, DNA regulation, lysis, and other essential functions as per the Prokaryotic Virus Orthologous Groups (pVOGs) database. Consequently, these could be classified as defective prophages (45). Plasmids can impart new traits to probiotic bacteria, including enhancements in bacterial metabolism, adherence, and even antibiotic resistance. However, if plasmids contribute antibiotic resistance to probiotic bacteria, this particular attribute is considered undesirable for a probiotic bacterium (46). However, in our study, we did not find any plasmid in the genomes of genus *Geobacillus*. We have also reported the same results previously in different studies (13, 27). The absence of mobile genetic elements, prophages, and plasmids reveals that the genomes of the genus *Geobacillus* are stable and good candidates as probiotics.

Bacteriocins, a diverse array of ribosomally synthesized antimicrobial peptides, provide probiotic bacteria with the advantageous ability to effectively combat other bacterial strains. They assist in the establishment of a producer bacterium within a specific environment, directly hinder the intrusion of competing strains or pathogens, and have the potential to alter the microbiota’s composition while influencing the host’s immune system (47). Certain studies have documented the production of bacteriocins within the Bifidobacterium genus (48). Similarly, several bacteriocins, including Circularin_A, ComX1, ComX4, Salivaricin_D, Sactipeptides, Pumilarin, and Geobacillin_I_like, were found across different genomes of *Geobacillus*. CRISPR-Cas systems are adaptive immune systems of microbes. These elements play a crucial role for bacteria in managing phage sequences found in the environment, particularly within the intricate ecosystem of the gut where a diverse viral community thrives. Phages can disrupt bacteria, significantly affecting the survival of bacterial populations, thereby impacting various parameters including the production of probiotics, fermentation duration, taste, and other essential factors (49). Given the resilience of phages to pasteurization and the difficulty in eliminating them, the pursuit of probiotics possessing the capacity to shield against phages and other genetic intruders like plasmids becomes an urgent necessity. In a previous study, 77% of 48 analyzed species of *Bifidobacterium* possess CRISPR-Cas systems (50). In our study, considering two prominent probiotic species of *Bifidobacterium* (*B. breve* and *B. longum*), we didn’t find any CRISPR-Cas-related genes. However, there were diverse genes related to CRISPR-Cas systems present in the genomes of *Geobacillus* such as *Cas1*, *Cas2*, *Cas3*, *Cas4*, *Cas5h*, *Cas6*, *Cas9*, *Cmr1*, *Cmr3*, *Cmr4*, *Cmr5*, and *Cmr6*. Thus, the presence of bacteriocins and CRISPR-Cas-related genes suggested the promising potential of probiotics among the genus *Geobacillus*.

A pivotal aspect of defining a probiotic strain is its stance on antibiotic resistance. Within the Qualified Presumption of Safety (QPS) criteria, evaluating probiotics involves examining the safety and antimicrobial resistance (51). Ideally, probiotics ought to be susceptible to at least two antibiotics or should lack inherent antimicrobial resistance (52). The genotypic approach involves full genome sequencing, including plasmids, to detect known antibiotic resistance (AR) genes. If these genes are surrounded by mobile elements or encoded within plasmids, it is advised against commercializing the strain. Safety considerations are crucial, especially if resistance involves clinically important antibiotics like vancomycin. If resistance affects less clinically relevant antibiotics, the risk is lower, but thorough evaluation is needed before commercialization (53). In our previous study (13) we found the genus *Geobacillus* is devoid of any antibiotic resistance. The results collide with the findings of Puopolo and co-workers, where *G. stearothermophilus* GF16 isolated from the active volcanic area was susceptible to different antibiotics used (even at the lowest concentrations of 5 µg/mL) (54). Likewise, the complete genome examination of G. thermoleovorans derived from geothermal springs indicates the lack of antimicrobial resistance genes (55). All of these findings of the absence of antibiotic susceptibility in *Geobacillus* in general (13). Similarly, in the present study, we only found two genes *vanY* and *vanT* related to vancomycin resistance. *VanY* acts as a D, D-carboxypeptidase, eliminating the terminal D-Ala from peptidoglycan to incorporate D-Lactate. The resulting D-Ala-D-Lac peptidoglycan subunits exhibit lower binding affinity with vancomycin in comparison to D-Ala-D-Ala. *VanY* serves as an “accessory” component in the Van cascade and isn’t an absolute necessity for vancomycin resistance (56). *Bifidobacterium* and *Lactobacillus* genomes were found to possess more genes apart from *vanY* and *vanT* such as *ImrD*, *rpsL*, and *qacJ*. A study has also reported the presence of erm, *ileS*, and *rpoB* genes in *B. breve*. Investigating the presence of virulence factors (including toxins and enzymes that could potentially heighten the microorganism’s pathogenicity) in the genome is crucial for identifying potential safety concerns. However, only a few toxin-like proteins and virulence factors are also absent in all the genomes of the genus *Geobacillus*. A few toxin-like proteins detected were *PhoH*, *MazF*, and *PemK* as compared to genus *Bifidobacterium* and *Lactobacillus* which possess an array of toxin-related genomes such as *FitB*, *PemK*, *MazF*, *ParB*, *RelB*, *RelE*, *MraZ*,*YoeB*, *PhoH*, and *HipA*. *PhoH2* proteins, found in various microorganisms across bacteria and archaea, comprise two domains: An N-terminal PIN-domain connected to a C-terminal *PhoH* domain. This fusion function as both an RNA helicase and ribonuclease. Mainly within mycobacteria, the genome encodes PhoH2 proteins alongside an adjacent gene, *phoAT*, believed to function as an antitoxin (57). The *mazEF* module, found on *Escherichia coli’s* chromosome, serves as a stress-induced toxin-antitoxin system that triggers programmed cell death in *E. coli*. Precisely, *mazF* encodes a resilient toxin, whereas *mazE* encodes a comparatively less stable antitoxin (58). In the *Bacillus anthracis* genome, a toxin-antitoxin (TA) module is present, consisting of *pemI* (antitoxin) and *pemK* (toxin). *PemK* functions as a powerful ribonuclease, demonstrating a preference for pyrimidines (specifically C and U), operating as a translational attenuator (59). Thus, in comparison to already known probiotic candidates of genus *Bifidobacterium* and *Lactobacillus,* the genus *Geobacillus* possess very few toxin-like protein coding genes. Thus, it has been proposed that genus *Geobacillus* may be good candidates with good probiotic potential.

Microbial genomes encode a multitude of genes responsible for Carbohydrate-active enzymes (CAZymes), whereas in humans, only 17 relevant ones have been identified (60). This discrepancy reveals that humans lack extensive enzyme machinery to utilize a wide array of complex carbohydrates. As a result, humans depend on a symbiotic co-metabolism with their microbiota to extract energy, particularly from carbohydrates that are otherwise indigestible (61). The dynamic CAZyme profile is influenced by factors like available carbohydrates, non-carbohydrate food sources, delivery methods, and individual lifestyles (62). Differences in host CAZymes or the lack of particular microbial species possessing distinct CAZymes can substantially impact how the host metabolizes various sources of carbohydrates (63), impacting gut microbiota metabolism and potentially affecting host health. Assessing microbial genetic diversity (CAZy-typing) helps predict which carbohydrates the host can metabolize and identifies underrepresented CAZyme families that might require supplementation through methods like microbiota transplantation or probiotics. In our study, we found an abundance of glycoside hydrolases followed by Glycosyl transferases and carbohydrate esterases among the genus *Geobacillus*. Thus, suggesting that they are capable of enzymatic degradation of various carbohydrates and consequently adding to the probiotic potential of these species. Thus, all the studied in silico features favor the probiotic nature of the genus *Geobacillus*. Yet, an assessment of *in vitro* characterization remains essential concerning probiotic attributes.

The unprecedented probiotic potential of the *Geobacillus* genus remains unexplored. With the Pan-Genome analysis’s advent, the intricate genetic makeup, the core genome concerning the essential function genes, and genes required to acclimatize in a new niche or secondary metabolite synthesis (conferred by the flexible genome) in a bacterium can be comprehensively disclosed. Analyzing the Pan-genome serves as a crucial method to provide insights into evolution and adaptation, and to explore potential probiotic characteristics in the group. Comparison to the genus *Lactobacillus* and *Bifidobacterium* in the current study revealed that the genus *Geobacillus* possesses an array of genes that aid in combatting stress conditions. Various genes conferring acidic tolerance and osmotic stress tolerance were found, with certain genes aiding in osmotic and oxidative stress tolerance confined to the genome of *Geobacillus* only. Determining the presence of mobile genetic elements (MGEs) remains paramount to instigating a strain with probiotic potential. Notably, *Geobacillus* appears stable, lacking mobile genetic elements (MGEs), antibiotic resistance genes, and mostly defective prophages. There were no observed plasmids, reducing the risk of horizontal gene transfer for antibiotic resistance. The presence of diverse bacteriocins and an abundance of CRISPR-Cas system genes in *Geobacillus* further suggests its potential against harmful elements like bacteriophages. Additionally, the prevalence of carbohydrate-active enzymes (CAZymes), especially glycoside hydrolases, glycosyl transferases, and carbohydrate esterases, hints at the ability of the genus *Geobacillus* to break down complex carbohydrates. This could enhance its potential as a probiotic, particularly in creating symbiotic food products when combined with prebiotics. While this study unveils promising probiotic traits in the genus *Geobacillus*, further in vitro and in vivo investigations are necessary to confirm these findings. Overall, the comparative genomic analysis sheds light on *Geobacillus*’ probiotic potential, yet empirical studies are imperative for substantiating these observations.

## Materials and Methods

### Genome Availability and Quality Control

All the 13 validly published *Geobacillus* species with correct names as per List of Prokaryotic names with Standing in Nomenclature (LPSN) and 5 randomly selected genomes (not validly published) of genus *Geobacillus* were selected and extracted in Fasta and GenBank formats from NCBI datasets https://www.ncbi.nlm.nih.gov/datasets/genome/. To access the probiotic potential of various studied *Geobacillus* species, 5 already known probiotic species such as *Lactobacillus acidophilus* ATCC 9224 [NZ_CP130437], *Lactobacillus delbrueckii* subsp. *lactis* DSM 20072 [CP022988], *Lactiplantibacillus plantarum* DSM 20174 [CP039121], *Bifidobacterium breve* DSM 20213 [BBAO00000000], and *Bifidobacterium longum* subsp. *infantis* ATCC 15697 [CP001095] were taken as control species. After the extraction of a total of 23 genomes, they were further analyzed for quality check, trimming, and alignment. The assessment of genome quality was done involved utilizing QUAST (version 4.4) (64). Additionally, to reconfirm genome completeness and identify any contamination, CheckM v1.0.18 (65) was employed, followed by trimming using the Trimmomatic program (version 0.36) (66). Multiple whole genome sequence alignments were conducted using Bowtie2 v2.3.2 (67). These analyses were performed on the KBase platform, available at https://www.kbase.us/ (68).

### Phylogenomic Analysis and Annotation

The study examined the phylogenetic relationships among *Geobacillus* species and their connections with other *Lactobacillus* and *Bifidobacterium* probiotic genomes used in this research, based on a defined set of 49 core, universally recognized genes from COG (Clusters of Orthologous Groups) gene families. This was performed by a KBase app Species Tree Builder v0.1.4 (68). This process begins by choosing a subset of publicly available KBase genomes that exhibit close relations to the genomes we’ve provided. Subsequently, these genomes undergo multiple sequence alignment (MSA) against 49 COG domains. The resulting alignments are meticulously refined using GBLOCKS to eliminate inadequately aligned sections within the MSA. These refined MSAs are concatenated, and a phylogenetic tree is generated using the maximum likelihood method via FastTree2 version 2.1.11. The phylogenomic analysis was also done by using the Codon Tree Test method of the phylogenetic tree-building service of Bacterial and Viral Bioinformatics Resource Center (BV-BRC) (https://www.bv-brc.org/) (69). The Codon Tree approach uses the RAxML program to analyze aligned proteins and coding DNA from single-copy genes after choosing single-copy BV-BRC PGFams. RaxML’s “Rapid” bootstrapping option was used to produce support values over 100 iterations.

The genomes were annotated using Prokka v1.14.5 (70) through the KBase platform. To assess the genes carrying known functions among the Prokka annotated genomes was done by the KBase app “View Function Profile for Genome-v1.4.0” and a heatmap was generated. It summarizes gene functions using canonical gene family assignment from COG, PFAM, TIGRFAM, and the SEED. Before this step, the domains of the Prokka annotated genomes were first annotated using the Domain Annotation app v1.0.10. (68).

### Detection of Genes Related to Probiotic Features

Genes involved in the mechanism of modulation of the immune system, vitamin biosynthesis, fatty acid synthesis, resistance to stress conditions (acid, bile, osmotic, and oxidative stress), adhesion, bacterial colonization, and molecular chaperones were detected from Prokka annotated genomes (70). These probiotic-related genes were compared with those of already known probiotic species considered in this study.

### Detection of Mobile Elements, Insertion Sequences, Plasmids, Prophages, Bacteriocins, and CRISPR-Cas Systems

The in-silico identification of plasmids among the studied genomes was done with PlasmidFinder 2.0 (71). Phage presence was assessed using Phigaro through the Proksee platform. (72). The mobile elements and insertion sequences were predicted by ICEFinder (73) and VRprofile2 (74). The putative bacteriocins were predicted by using BAGEL4 (75), and the presence of Clustered Regularly Interspaced Short Palindromic Repeats (CRISPRs) and Cas proteins were analyzed with the CRISPRCasfinder v4.2.20 tool (76) using Proksee (72) platform and were also cross-checked in Prokka annotations of genomes using KBase platform.

### Detection of Antibiotic Resistance Genes, Virulence Factors, and Toxins

The assessment of antibiotic resistance gene predictions was conducted using ABRIcate v1.0.1 within the Galaxy platform, RGI (Resistance Gene Identifier) 6.0.3, and CARD (Comprehensive Antibiotic Resistance Database) (77) using PATRIC (BV-BRC v3.32.13a) and Proksee platforms. The virulence factors were predicted with PATRIC (BV-BRC v3.32.13a) and ABRIcate v1.0.1 against VFDB (virulence factor data base) (69). The toxins were predicted with TAser (https://shiny.bioinformatics.unibe.ch/apps/taser/) using the TASmania database (78). Only those genes were considered with hmm_E_value less than 1e-15.

### Identification of Carbohydrate-active Enzymes (CAZyme)

The carbohydrate-active enzymes were identified by annotating the genome sequences through dbCAN (79) with dbCAN2 HMMs of CAZy families - v10 app in KBase. This method scans protein sequences found in Genomes using a set of Hidden Markov Models (HMMs) from the dbCAN2 CAZy family collection or CAZy database (http://www.cazy.org/). It uses HMMER software v3.3.2 is installed from http://hmmer.org.

### Pangenome Analysis

The Pangenome analysis was performed with Genome Comparison SDK v.0.0.7 app in KBase. The KBase algorithm for computing the protein pangenome relies on a k-mer-based approach, providing significant advantages. It entails identifying uniquely distinct k-mers, specifically of length 8, present in each protein across all genomes. Phylogenetic Pangenome Accumulation v1.4.0 (https://kbase.us/applist/apps/kb_phylogenomics/view_pan_phylo/) was run to view the pangenome in phylogenetic context and to compare among two genome sets (between *Geobacillus* and *Lactobacillus/Bifidobacterium*). This application allows to dissecting of pangenome categories using a species tree to determine the entry and exit of gene families in the branch of interest.

The Pangenome analysis was also done with IPGA (Integrated Prokaryotes Genome and Pan-Genome Analysis) v1.09 (80). Before any analysis quality control is done on input genomes. After QC, the genes of all filtered genomes are predicted using prokka. Using up to eight different types of software such as OrthoMCL, PanOCT, Roary, OrthoFinder, panX, Panaroo, PPanGGoLiN, and PEPPAN each gene is annotated against the COG database to provide a unique pan-genome profile. To assist users in choosing the optimal pan-genome profile from the possibly inconsistent findings, IPGA extracts all orthologous gene clusters and then assigns a score to each of them (80). Pangenome Circle Plot was formed using Pangenome Circle Plot-v1.2.0 in KBase (https://kbase.us/applist/apps/kb_phylogenomics/view_pan_circle_plot/release). This app allows the overlapping membership of genes against a base genome.

## Acknowledgments

Dr. Ishfaq Nabi Najar would like to thank DBT-RA, Department of Biotechnology, Govt. of India, for providing the DBT-RA Fellowship (**DBT-RA/2022/JULY/N/2877**) for research work. I would also like to thank CSIR IIIM Jammu, for providing me lab space to carry out my research.

## Ethics approval

Not applicable

## Funding

Not applicable.

## Credit authorship contribution statement

**Ishfaq Nabi Najar:** Conceptualization, Data Curation, Analysis, Writing – original draft. **Prayatna Sharma:** Analysis and Writing– original draft. **Rohit Das:** Analysis and Editing – original draft. **Krishnendu Mondal, Ashish Kumar Singh**, **Anu Radha**, **Varsha Sharma**, **Sonali Sharma**-Data Curation, Writing and Editing – original draft, **Nagendra Thakur**: Review & Editing. **Sumit G Gandhi**: Review & Editing. **Vinod Kumar:** Conceptualization, Review & Editing.

## Declaration of Competing Interest

The authors declare no competing interests.

